# A novel analysis pipeline and experimental design for measuring expanded repeat instability

**DOI:** 10.64898/2026.03.12.707943

**Authors:** Christopher Smith, Ruban Rex Peter Durairaj, Emma L. Randall, Alys N. Aston, Laura Heraty, Waheba Elsayed, Alvaro Murillo, Vincent Dion

## Abstract

The expansion of short tandem repeats is a feature of over 60 different human diseases. Ongoing somatic instability throughout a patient’s lifetime can influence disease progression and has emerged as a therapeutic target. Understanding its mechanism is essential for the identification of both drug targets and therapeutic interventions. A major obstacle towards this translational goal has been to measure changes in repeat size distributions. To address this we provide a new analysis method and accompanying software that generates delta plots, extracts the instability frequency from targeted long-read sequencing data, the instability bias, and the average size of the changes. It further provides statistical analysis for comparison between treatments. We show its applicability to non-dividing cells and *in vivo* datasets. Moreover, we have developed a streamlined experimental design for dividing cells, Single Clone-based Instability Assay (SCIA), that saves weeks in assessing the effect of a gene knockout on repeat instability and is ideal for an initial screen. We have validated the approach using *FAN1, PMS1,* and *MLH1* knockouts in cell lines and induced pluripotent stem cells. We find that *FAN1* knockout clones showed, as expected, an increased frequency of repeat expansion but the size of the expansions were smaller. This highlights the wealth of information that can be extracted and the potential for novel insights into the mechanism of repeat instability.

## Background

Expanded short tandem repeats are genetically unstable, leading to somatic mosaicism within a tissue over time. This phenomenon of somatic instability has now been observed in most, if not all, of the over 60 diseases caused by the expansion of short tandem repeats (Chen et al. 2024; Khristich and Mirkin 2020; López Castel et al. 2010). Somatic instability has a central role in the aetiology of expanded repeat disorders (Bettencourt et al. 2016; Donaldson et al. 2025; Genetic Modifiers of Huntington’s Disease (GeM-HD) Consortium 2019) and has emerged as a major target for therapeutic intervention, at least for Huntington’s disease (Donaldson et al. 2025; Binda et al. 2024; Marzec et al. 2025).

Expanded CAG/CTG repeats are the best studied model of repeat instability. Repeats beyond 34 units in length form stable non-B DNA structures containing mismatched hairpins (Mirkin 2007). These structures are then recognized by mismatch repair complexes, which nick the DNA and create a substrate for error-prone DNA synthesis leading to repeat expansion (Iyer and Pluciennik 2021; Fernandez et al. 2026). The MutLγ complex, composed of MLH1 and MLH3, is thought to provide that first nick opposite a CAG or CTG hairpin (Iyer and Pluciennik 2021; Senoussi et al. 2025), whereas FAN1, an endonuclease that binds MLH1 and prevents repeat expansion, is thought to counter this process by physically antagonising the activity of MutLγ and by digesting the hairpin strand (Senoussi et al. 2025; Deshmukh et al. 2021a; Li et al. 2025; Phadte et al. 2023). As a consequence, FAN1 prevents repeat expansion in multiple cellular and *in vivo* models (Goold et al. 2019, 2021; Loupe et al. 2020; Mouro Pinto et al. 2025; Porro et al. 2021; Zhao and Usdin 2018).

However, the molecular mechanisms of repeat instability are incompletely understood. Indeed, 40 modifiers of CAG/CTG repeat instability have been identified in mammalian systems but how they are working together in the pathway(s) that cause(s) repeat instability and their genetic interactions with mismatch repair proteins is not clear.

A major obstacle towards understanding the mechanism of repeat instability and testing the effect of potential genetic targets is measuring changes in repeat size mosaicism. This is in part because repeat size distribution within a sample is complex with potentially different mechanisms leading to small and large changes in repeat size (Kim et al. 2017). Moreover, changes brought about by a treatment can affect the bias towards contractions or expansions, or alter the rate of instability. There is currently no analysis pipeline that captures the whole spectrum of the potential changes. Instead, the most widely used measure, the instability index (Lee et al. 2010), amalgamates both the size and the frequency of the changes in a single number. It masks the potential underlying mechanisms and reduces the statistical power of the analysis.

Here we provide an end-to-end solution for measuring repeat size changes based on targeted long-read sequencing. We developed a graphical user interface and the accompanying statistical tests to make the pipeline widely accessible. In addition to applying the approach to a wide array of datasets, we optimised a novel and robust experimental system that can save weeks when uncovering the effect of a genetic mutation and avoiding long-term bulk cultures and the need for reporter assays. We further show that novel mechanistic insights can be gained using this approach.

## Results

### Quantification of repeat size mosaicism in multiple cell types

One straightforward way of quantifying repeat size distributions is to use the instability index (Lee et al. 2010). This is a weighted measurement of repeat size normalized to the mode of the starting or untreated population. Instability indices, however, have some limitations. Specifically, they bundle together repeat size changes and the frequency of those changes, leading to an incomplete understanding of the repeat instability mechanism. To extract as much of the data as possible, we opted to present changes in repeat size distributions as delta plots. This works by using targeted sequencing to determine repeat size distributions for a control and a treated population and subtracting one profile from the other (Figure 1a). We then plot that difference, or delta, resulting in graphs that highlight the changes in the repeat size independently of the shape of the starting distribution. From these graphs, we can obtain a measure of the frequency of instability (i.e., the proportion of the alleles that have changed in size), the bias ratio of expansions to contractions, and the average size of the changes (Figure 1b).

**Figure 1:**
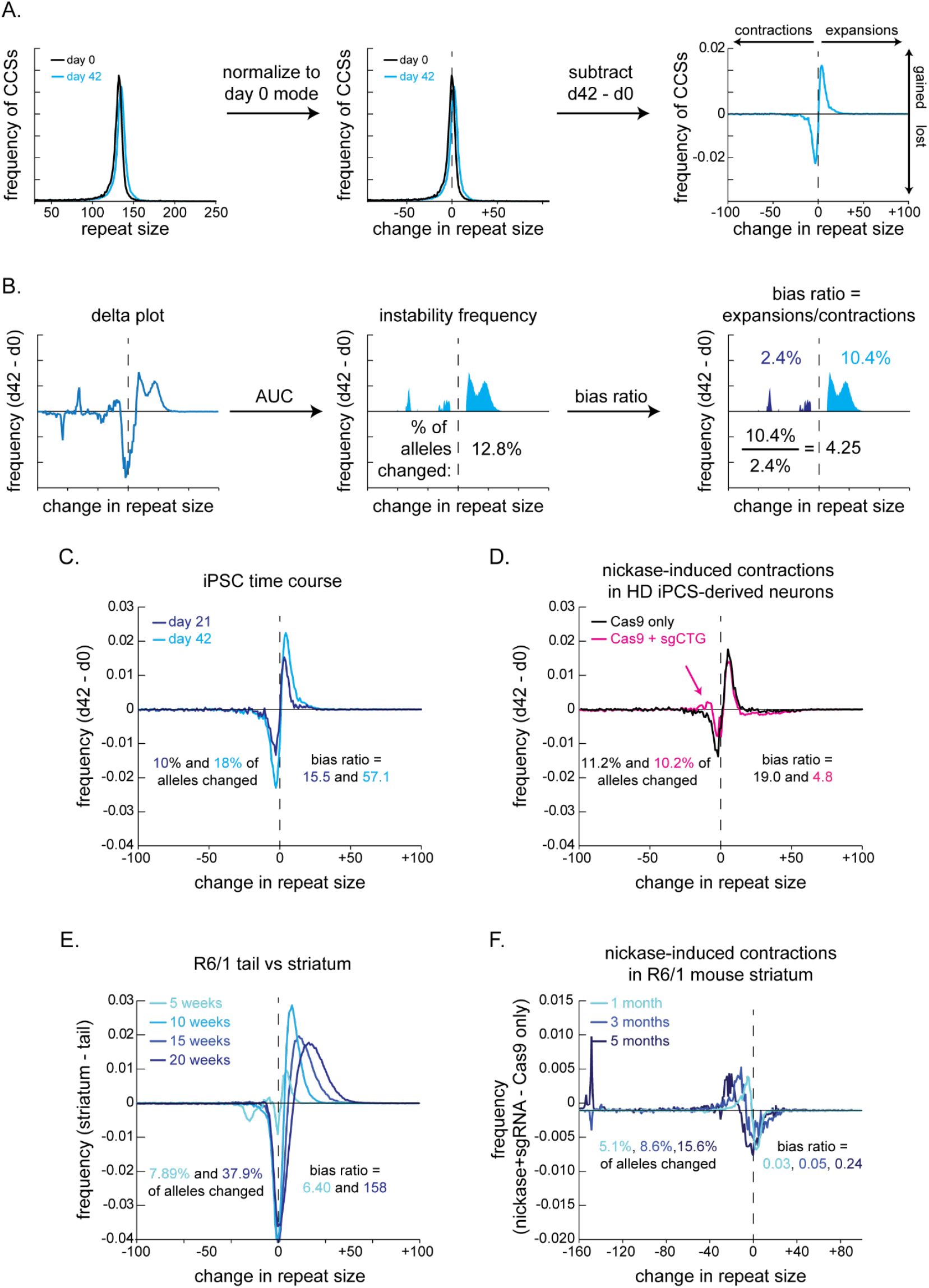
Delta plots are widely applicable to visualise changes in repeat size. A) Method for generating delta plots. The histograms of repeat size distributions for a treated (in this case iPSC from HD at the start of the experiment (day 0, black) and treated (in this case grown for 42 days, light blue) are adjusted so that the mode of the culture at day 0 is taken as no change in repeat size. Then the two distributions are subtracted to generate delta plots showing changes in repeat size around the mode. B) Instability frequency (based on the area under the curve (AUC) and the bias ratios (of expansions over contractions) can then be calculated. Example shown is from GFP(CAG)116 cells that have been grown for 42 days as single clones (see below). C) Delta plots of iPSCs grown over 21 and 42 days. D) HD iPSC-derived neurons transduced with a lentivirus expressing either Cas9D10A alone or Cas9D10A+sgCTG. This latter condition is known to induce a modest number of contractions (Murillo et al. 2025). E) Delta plots can be used to quantify the amount of repeat expansion accumulating over time in the R6/1 striatum compared to the more stable tail. F) Delta plots of R6/1 striata comparing hemispheres injected with Cas9D10A alone or Cas9D10A+sgCTG to induce contractions. The mice were injected with AAVs 2 days after birth and followed over 5 months. The data is from (Murillo et al. 2025).

We have applied this approach to a time-course experiment where we used HD- derived induced pluripotent stem cells (iPSCs). These cells contain two alleles, an expanded at 135 and a non-expanded allele at 19 CAGs at the *HTT* locus (Murillo et al. 2025; HD iPSC Consortium 2012). We isolated DNA from the starting population (day 0) and of the population over time (days 21 and 42) and performed long-read sequencing. For this, we opted for PacBio Single Molecule Real Time (SMRT) sequencing because it is less error prone than Oxford Nanopore Sequencing and accommodates a greater range of repeat sizes than MiSeq sequencing (Taylor et al. 2022). We used a PCR-based library preparation for convenience and lower costs (Murillo et al. 2025; Larin et al. 2024) and ran Repeat Detector (Taylor et al. 2022) to determine repeat size distributions. We found that the delta plots highlighted the expansions that occurred in these lines over time with an average instability frequency at 21 days of 10.0%, a bias ratio of 15.5 expansions for every contraction, and an average expansion size of 6.2 ± 9 (Figure 1c, Table 1). Three weeks later, the instability frequency increased to 18% and the bias ratio to 57.1, indicative of an accumulation of expansions (Figure 1c, Table 1), as expected. We also applied this pipeline to HD-iPSC-derived neurons and compared those treated with the Cas9D10A nickase with or without a sgRNA against the repeat tract so as to contract them ([26], Figure 1d, Table 1), and to mouse brain samples over time compared to the more stable tail tissue (Figure 1e, Table 1) or treated or not with the Cas9 nickase to induce contractions ((Murillo et al. 2025), Figure 1f, Table 1). We conclude that the analysis pipeline is widely applicable to datasets from multiple experimental systems, including non-dividing cells and brain tissues.

**Table 1:**
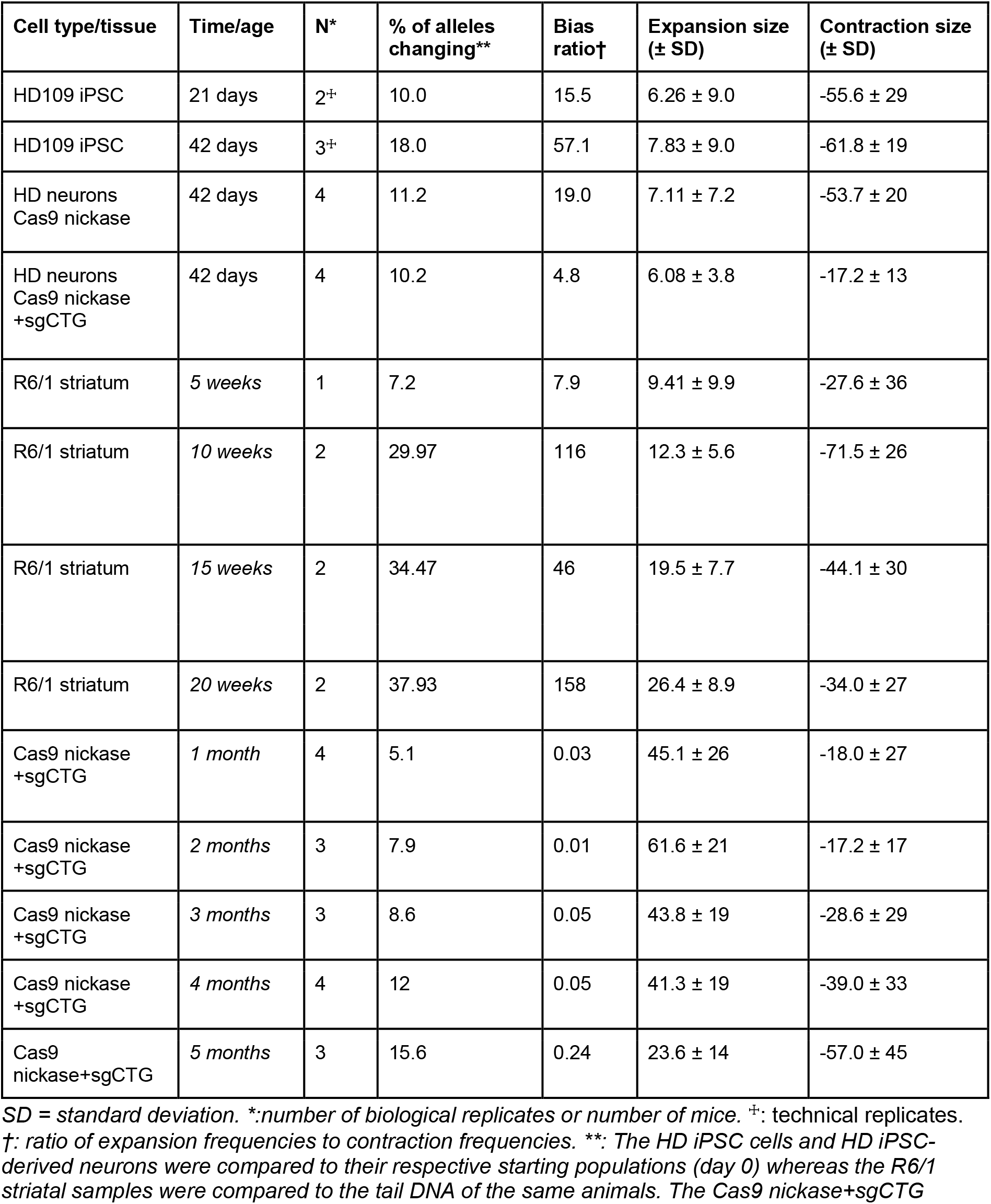
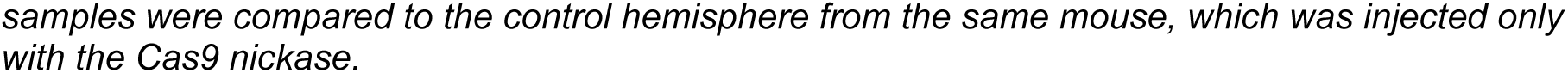
Instability statistics for HD iPSCs, HD iPSC-derived neurons, and striatal samples.

| Cell type/tissue | Time/age | N* | % of alleles changing** | Bias ratio† | Expansion size (± SD) | Contraction size (± SD) |
| --- | --- | --- | --- | --- | --- | --- |
| HD109 iPSC | 21 days | 2<sup>±</sup> | 10.0 | 15.5 | 6.26 ± 9.0 | -55.6 ± 29 |
| HD109 iPSC | 42 days | 3<sup>±</sup> | 18.0 | 57.1 | 7.83 ± 9.0 | -61.8 ± 19 |
| HD neurons<br>Cas9 nickase | 42 days | 4 | 11.2 | 19.0 | 7.11 ± 7.2 | -53.7 ± 20 |
| HD neurons<br>Cas9 nickase<br>+sgCTG | 42 days | 4 | 10.2 | 4.8 | 6.08 ± 3.8 | -17.2 ± 13 |
| R6/1 striatum | 5 weeks | 1 | 7.2 | 7.9 | 9.41 ± 9.9 | -27.6 ± 36 |
| R6/1 striatum | 10 weeks | 2 | 29.97 | 116 | 12.3 ± 5.6 | -71.5 ± 26 |
| R6/1 striatum | 15 weeks | 2 | 34.47 | 46 | 19.5 ± 7.7 | -44.1 ± 30 |
| R6/1 striatum | 20 weeks | 2 | 37.93 | 158 | 26.4 ± 8.9 | -34.0 ± 27 |
| Cas9 nickase<br>+sgCTG | 1 month | 4 | 5.1 | 0.03 | 45.1 ± 26 | -18.0 ± 27 |
| Cas9 nickase<br>+sgCTG | 2 months | 3 | 7.9 | 0.01 | 61.6 ± 21 | -17.2 ± 17 |
| Cas9 nickase<br>+sgCTG | 3 months | 3 | 8.6 | 0.05 | 43.8 ± 19 | -28.6 ± 29 |
| Cas9 nickase<br>+sgCTG | 4 months | 4 | 12 | 0.05 | 41.3 ± 19 | -39.0 ± 33 |
| Cas9<br>nickase+sgCTG | 5 months | 3 | 15.6 | 0.24 | 23.6 ± 14 | -57.0 ± 45 |

SD = standard deviation. \*:number of biological replicates or number of mice. <sup>±</sup>: technical replicates. †: ratio of expansion frequencies to contraction frequencies. \*\*: The HD iPSC cells and HD iPSC-derived neurons were compared to their respective starting populations (day 0) whereas the R6/1 striatal samples were compared to the tail DNA of the same animals. The Cas9 nickase+sgCTG
*samples were compared to the control hemisphere from the same mouse, which was injected only with the Cas9 nickase.*

### A software for the analysis of changes in repeat size distributions

To promote the use of delta plots and of the analysis pipeline, we have developed software with a graphical user interface (GUI) (Supplementary Video 1, Supplementary Figure 1). It requires a docker container, the docker object .tar file, and the FASTA files containing the circular consensus sequences (CCSs) or reads as input. The software runs Repeat Detector (Taylor et al. 2022), which determines the size of the repeat for each CCS and outputs text files and plots the histograms of frequencies against repeat sizes. There is an option to calculate the instability index. The second module calculates delta plots from the normalised histogram data. Two independent datasets, each containing both control and treatment histograms, can be loaded simultaneously. It determines the area under the curve, the average size of the expansions and of the contractions, and the bias ratio. There is an option to perform a KS test to compare two genotypes. Of note, the software is not limited to SMRT sequencing data and can be used with an Illumina MiSeq or, with some modifications to the Repeat Detector profile, Oxford Nanopore Technologies. One limitation of the current version is that some experimental designs, for example when we compared experimental replicates, each with their own controls and treated samples (e.g., comparing two mouse hemispheres, one treated, the other not), the software will extract all the information for the individual experimental replicates, but the averaging needs to be done by hand. The software is freely available via https://github.com/DionLab/SCIA, with the version controlled code found at https://doi.org/10.5281/zenodo.18850398.

### Minimising the effect of clonal expansion in bulk culturing

Immortalised cells that rapidly divide have been extensively used for studying repeat expansion (Dion 2014; Massey et al. 2018). This is because they are easy and cheap to culture, and are genetically manipulated at will. Importantly, they have genetic requirements for repeat expansion that are similar to those of non-dividing systems, including the reliance on mismatch repair for driving expansions and FAN1 in limiting it (Pluciennik et al. 2013; Goold et al. 2021, 2019; Porro et al. 2021; Zhao and Usdin 2018; Wheeler and Dion 2021; Massey et al. 2018; Donaldson et al. 2025). They are a screening tool where the results need to be confirmed in slower, more tedious, and more expansive disease models. Here we focus on speeding up the initial screening step.

Repeat expansion occurs at a rate of only a few repeats per week and thus measuring changes often requires long-term culturing. This puts a selection pressure on the culture such that cells that grow faster take over the population, irrespective of their repeat sizes, leading to inaccurate results. We have observed this in one culture of HEK293-derived GFP(CAG)_89_ cells (Figure 2). These cells were characterized previously and contain an ectopic CAG repeat inserted on chromosome 12. Transcription through the repeat is driven by a doxycycline (dox) -inducible promoter (Cinesi et al. 2016; Ruiz Buendía et al. 2020; Santillan et al. 2014). This particular cell population showed two main repeat sizes, as measured by PCR, in which a shorter repeat took over the population within the course of a month (Figure 2ab). This is unlikely due to a selective advantage provided by the short repeat tract because we had not seen this in other cultures of this line before (e.g.,(Yang et al. 2022)) and making single-cell isolates and culturing them for 21 days did not lead to the same accumulation of short repeats (Figure 2c). These results support the idea that bulk populations cultured for an extended period can lead to artefacts and should be avoided if possible.

**Figure 2:**
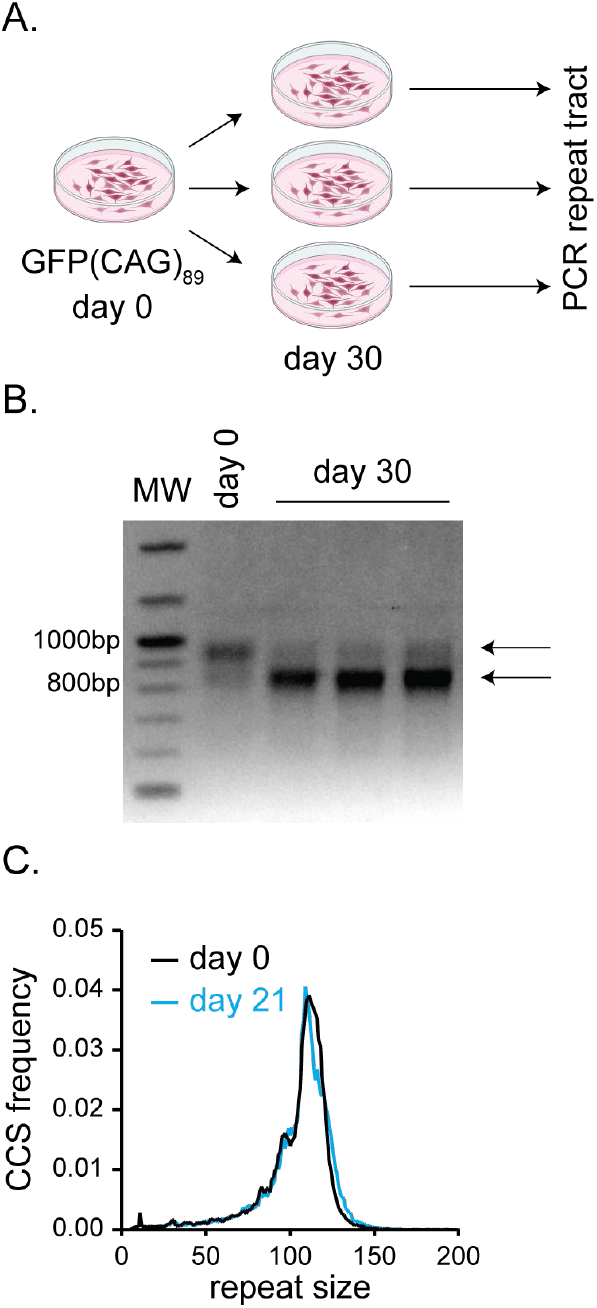
Artefact in measuring repeat instability can result from bulk culturing. A) Timeline for bulk culturing of GFP(CAG)_89_ cells. B) PCR products of the experiments done in A. Arrows show the two main bands for repeat sizes in these cultures. C) Single-cell isolate from GFP(CAG)_89_ with a repeat size of 111 does not show the accumulation of a shorter band seen in parental culture shown in B.

### Single Clone-based Instability Assay (SCIA)

The classical way of preventing artefacts due to bulk culturing when measuring rates of mutation is to use a fluctuation analysis (Lea and Coulson 1949). In this experimental design, multiple single cell clones are produced from a starting population to assay the rates of mutations. In the context of expanded repeats, fluctuation analyses are used extensively in dividing yeast cells (Polleys et al. 2017; Radchenko et al. 2018) but have been used only once, to our knowledge, in mammalian cells (Lin et al. 2006). The advantage of this experimental design is that each clone grows at their own rate and will not be in competition with the others, reducing the possibility that one mutant cell takes over the culture. One reason why this approach has been scarcely used in cultured mammalian cells, however, is that it requires a selectable reporter to detect changes in repeat size, which measures only specific types of instability events.

We sought to provide a selection-free experimental design that is widely applicable. To this end, we developed SCIA to test the hypothesis that using single clones coupled to long-read sequencing solves the issues of bulk culturing and nullifies the need for a reporter system. Having single clones also allows us to increase the number of replicates as most of the work is hands off, waiting for the clones to grow. We found that sorting up to 12 individual clones from a starting population was easily manageable and provided robust statistical analyses compared to the usual 3 or 4 replicates.

We sorted individual GFP(CAG)_116_ cells from an initial population (day 0) and grew them independently over 42 days (Figure 3a). We found that individual clones often showed narrower distributions of repeat sizes compared to the starting population, presumably because there has not been as much time for changes in repeat size to accumulate in the clones compared to the starting population (Figure 3b). A few clones had more than one peak, presumably because a large change in repeat size occurred shortly after the single cells were plated (e.g., clone 11). We excluded cases where two colonies were found in a single well after single-cell plating. We compared each single clone cultured for 42 days to the distribution from which they came at day 0 (Figure 3c). For these cells, the delta plot shows that expansions have increased in frequency at the expense of shorter repeats, which decrease in frequency over the time period (bias ratio of expansion frequency over contraction frequency = 4.25 - Figure 3c). Overall, 12.8% of the alleles changed in size as judged by the area under the curve. The average size changes were 25.7 ± 11 CAGs for expansions and -39.6 ± 24 for contractions (Figure 3c, Table 1). Overall, the SCIA pipeline provides a more complete view of repeat instability compared to instability indices.

**Figure 3:**
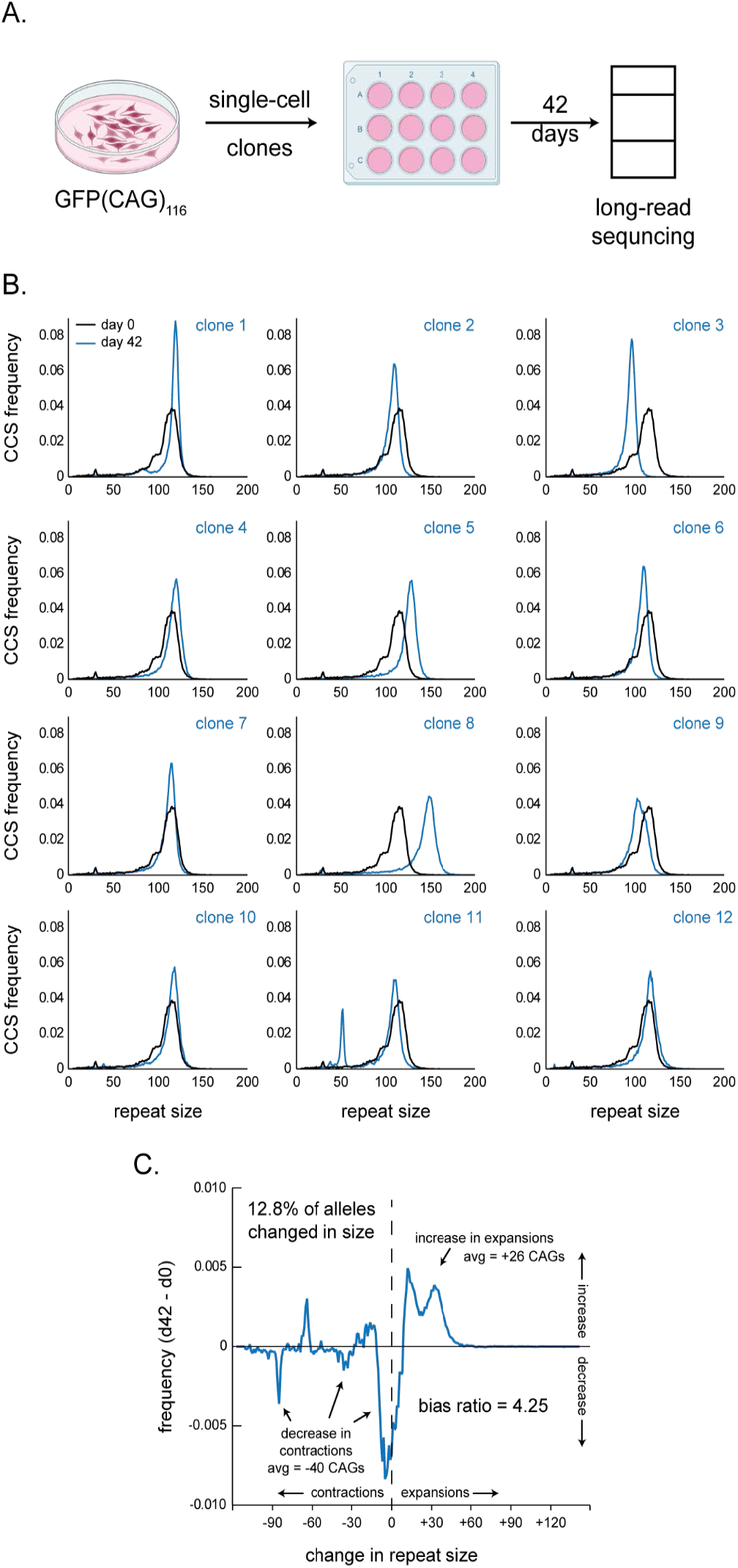
Single clone-based instability assay. A) Individual clones are grown from single cells isolated from a bulk population. After growing up the clones, the DNA is isolated and sequenced using targeted long-read Single molecule real-time (SMRT) sequencing. The repeat size is then compared to the bulk population (day 0). B) Histograms of repeat size distribution for each single clone compared to the starting population. C) Delta plot of GFP(CAG)_116_ between the starting population at day 0 and the average of the 12 clones seen in B at day 42. The percentage of alleles that changed in size is derived from the area under the curve and the average size (avg) of expansions and contractions are weighted averages. The bias ratio is the area under the curve for expansions divided by that of contractions.

### Reproducing the effect of repeat size on repeat instability

Next we sought to validate the experimental design by confirming the well-known effect of repeat size on repeat expansion rates, with longer repeats being more unstable and leading to more expansions (Veitch et al. 2007). The delta plots revealed a statistically significant difference between the two lines (Figure 4a, two-tailed KS test, P=0.0005). Also, as expected, the line with the larger repeat size had a higher bias ratio (1.48 expansion for every contraction in GFP(CAG)_81_ compared to 4.25 in GFP(CAG)_116_). The size of the changes were nearly four times larger in the line with the longer repeat tract. Surprisingly, the frequencies of the changes were slightly lower in the line with the larger repeat size (16.9% vs 12.8% for 81 repeats and 116 repeats, respectively - Table 2). We conclude that our new experimental design can detect differences in repeat size dynamics between two lines with different repeat sizes and uncover richer instability patterns than were previously possible.

**Figure 4:**
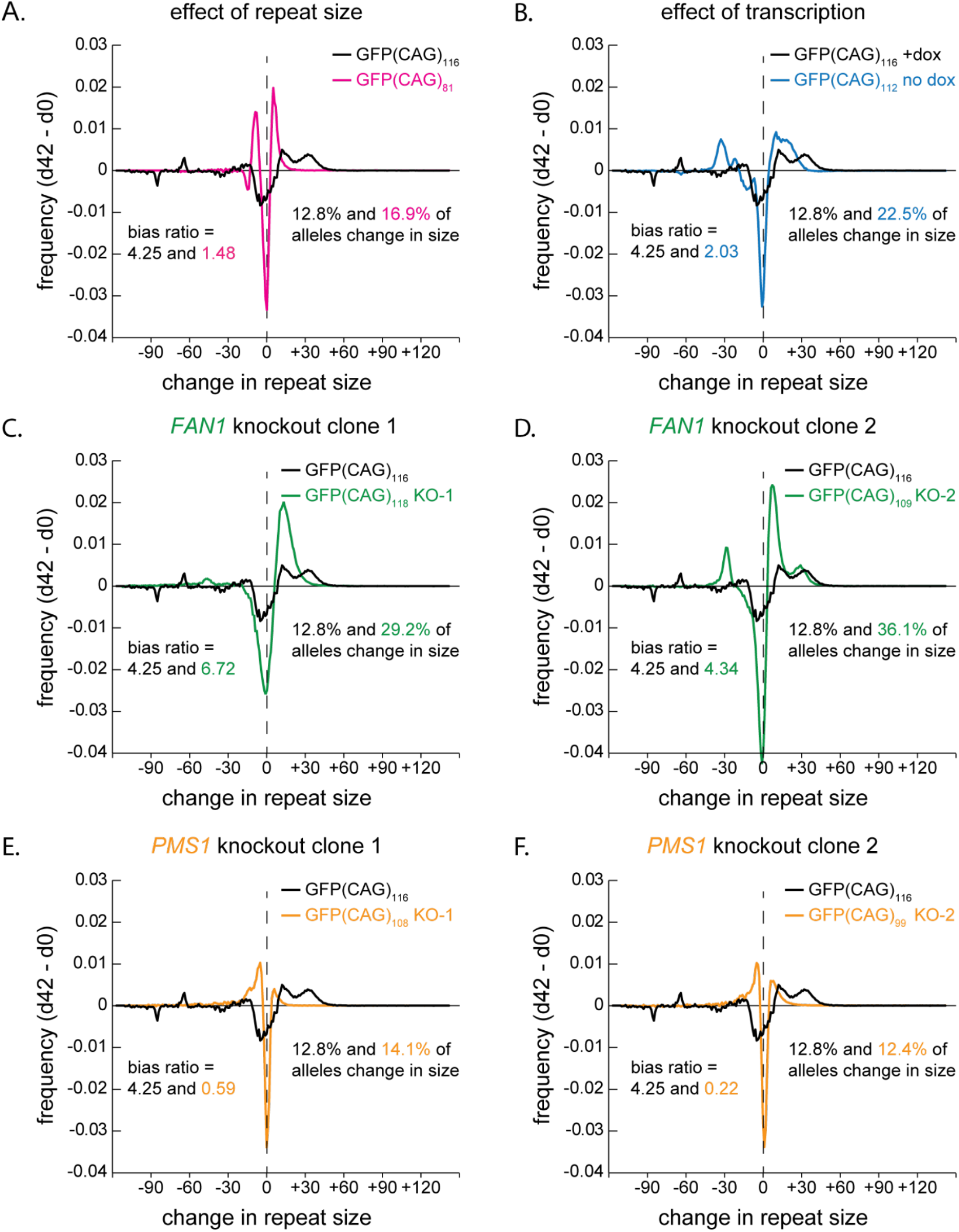
Validation of SCIA using genetic and pharmacological treatment. A) Delta plots showing the different in repeat instability over 42 days comparing a line with 116 repeats (black) and one with 81 repeats (pink) (transcription on). B) Delta plots showing that transcription (+dox, black) affects repeat instability compared to uninduced transcription (-dox, blue). C and D) Delta plots comparing two FAN1 KO clones (green) to a parental cell line (black). The experiments were done in the presence of dox. E and F) Delta plots comparing two PMS1 KO clones (orange) to its parental cell line (black). The experiments were done in the presence of dox. The data from GFP(CAG)_116_ with dox (black) has been used in every panel. We found that the distribution was similar between experiments and we therefore averaged them together.

**Table 2:** Instability statistics determined by the SCIA pipeline.

| Cell type | Repeat size | Genotype | Treatment | No. of clones | % of alleles changing | Bias ratio† | Expansion size (± SD) | Contraction size (± SD) | KS test P-value |
| --- | --- | --- | --- | --- | --- | --- | --- | --- | --- |
| GFP(CAG)<sub>x</sub> | 116 | WT | +dox | 12 | 12.8% | 4.25 | 25.7 ± 11 | -39.6 ± 24 | 0.0017 § |
| GFP(CAG)<sub>x</sub> | 81 | WT | +dox | 12 | 16.9% | 1.48 | 7.3 ± 4.6 | -10.0 ± 5.4 | 0.0005 * |
| GFP(CAG)<sub>x</sub> | 110 | WT | - | 23 | 22.5% | 2.03 | 16.3 ± 6.9 | -31.5 ± 6.5 | 0.0017 * |
| GFP(CAG)<sub>x</sub> | 118 | *FAN1* KO-1 | +dox | 11 | 29.2% | 6.72 | 16.3 ± 7.8 | -52.7 ± 20 | <0.0001 * |
| GFP(CAG)<sub>x</sub> | 109 | *FAN1* KO-2 | +dox | 12 | 36.1% | 4.34 | 14.1 ± 9.9 | -30.3 ± 5.4 | 0.0087 * |
| GFP(CAG)<sub>x</sub> | 108 | *PMS1* KO-1 | +dox | 7 | 14.1% | 0.59 | 10.8 ± 8.8 | -12.9 ± 11 | <0.0001 * |
| GFP(CAG)<sub>x</sub> | 99 | *PMS1* KO-2 | +dox | 11 | 12.4% | 0.22 | 9.1 ± 9.0 | -15.9 ± 16 | <0.0001 * |
| GFP(CAG)<sub>x</sub> | 109 | *MLH1* KO | - | 9 | 27.0% | 0.08 | 21.4 ± 4.4 | -9.8 ± 8.9 | <0.0001§ |
| HD iPSCs | 134 | *MLH1*<sup>+/-</sup> | - | 7 | 19.2% | 86.2 | 10.8 ± 5.8 | -87.1 ± 4.7 | N/A |
| HD iPSCs | 133 | *MLH1*<sup>-/-</sup> | - | 14 | 7.41% | 0.83 | 4.23 ± 7.5 | -6.79 ± 12 | 0.0004 ** |

WT = parental wild-type. dox = doxycycline. SD = standard deviation. KS = Kolmogorov-Smirnov comparing the delta plot distributions. §: compared to GFP(CAG)<sub>110</sub> (no dox), \*: compared to GFP(CAG)<sub>116</sub> (+dox), †: calculated from the area under the curve with expansions being increased compared to the mode at day 0 and contractions being decreased in repeat size compared to day 0. \*\*: comparing HD iPSCs *MLH1*<sup>-/-</sup> to HD iPSCs *MLH1*<sup>+/-</sup>.

### Probing the effect of transcription on repeat instability with SCIA

Transcription is known to destabilize repeat tracts (Lin et al. 2006, 2009; Mathews et al. 2025; Nakamori et al. 2011). The mechanism is thought to involve mismatch repair (Lin et al. 2006), transcription-coupled repair (Lin et al. 2009) and/or R-loop processing (Polleys et al. 2017; Lin et al. 2010; Reddy et al. 2011). However, one recent study has suggested that in RPE-1 derived cells, transcription reduces repeat expansion rates through an unknown mechanism (McLean et al. 2024). We showed little difference in expansions accumulating when transcription is allowed through the repeat tract in HEK293-derived cells using small-pool PCR (Yang et al. 2022). The differences may be because of the cell types used (Dion 2014), the location of the repeat compared to origin of replication (Cleary and Pearson 2005), whether contractions or expansions are being monitored, or the assay used to detect changes, among other considerations.

Here we tested the role of transcription by taking advantage of the dox-inducible promoter found in the GFP(CAG)_116_ cells (Cinesi et al. 2016; Santillan et al. 2014). We repeated a SCIA assay without the addition of dox (Figure 4b). We found that transcription significantly shifted the instability curves (KS test, P = 0.0017), but in ways that were not completely expected. For instance, the addition of dox reduced the instability frequency from 22.5% to 12.8% but increased the size of the expansions and the bias ratio, implying a larger expansion bias and an increased size of the resulting alleles. The latter is in line with a previous study with this line [36], supporting the idea that the SCIA protocol is robust.

### Validation of SCIA using genetic knockout lines

Next we determined whether known genetic modifiers of repeat instability could be tested using SCIA. One particularly relevant factor is FAN1, an endonuclease that can process secondary structures containing repeats and prevent repeat expansion (Deshmukh et al. 2021b). We hypothesised that knocking out *FAN1* in our GFP(CAG)_116_ cells would increase the frequency of repeat expansion. To do so, we made two independent clones that were knocked out for *FAN1,* confirmed the deletion by PCR and Sanger sequencing, and found no FAN1 protein by western blot (Supplementary Figure 2). When we compared changes in repeat size mosaicism, we found that both *FAN1* knockout lines were significantly different from the parental line (P ≦ 0.0087). Both clones showed an increase in the instability frequency (29.2% and 36.1% compared to 12.8% in the parent line), and a decrease in the size of the expansions (16.3 CAGs and 14.1 CAGs on average compared to 25.7 CAGs in the parent line - Figure 4cd, Table 1). These results confirm what has been seen in other systems: that FAN1 inhibits repeat expansion, but adds more information by suggesting that size of the expansions is smaller, but more frequent than in wild type cells.

We then knocked out *PMS1*, which is known to promote repeat expansion (Mouro Pinto et al. 2025; McLean et al. 2024; Ferguson et al. 2024; Hayward et al. 2024; Miller et al. 2020; Wang et al. 2025). We made two independent knockout lines (Supplementary Figure 3) using CRISPR and tested the lines with SCIA. Both *PMS1* knockout clones showed instability patterns significantly different from the parental line (P<0.0001). We also found that the instability frequency had not changed, but that the bias ratio dropped from 4.25 in the parental line to below 1 in the clones, implying a bias towards contractions in the knockout lines (Figure 4ef, Table 1). These results argue that, as in other systems, *PMS1* deletion in this line leads to a lower level of repeat expansion.

### A quick protocol to evaluate the effect of a gene on repeat instability

To overcome a major issue in evaluating the effect of a gene on repeat instability, time, we took advantage of the single-clone approach and treated each clone as an independent experiment. Classically, one would generate gene knockouts and screen clones for repeat size, protein expression, genetic mutation and perform functional assays. Then, two or more clones are selected and fed into a repeat instability assay. The shortcut implemented here (Figure 5a) is to transfect a bulk culture with the Cas9 RNPs and feed that transfected population directly into the SCIA pipeline. Validation of knockout clones is done in parallel as the colonies grow over the 42 day-long assay. This bypasses the need to first make and characterize the mutant clones. If needed, individual clones are available at the end of the experiment for further characterization.

**Figure 5:**
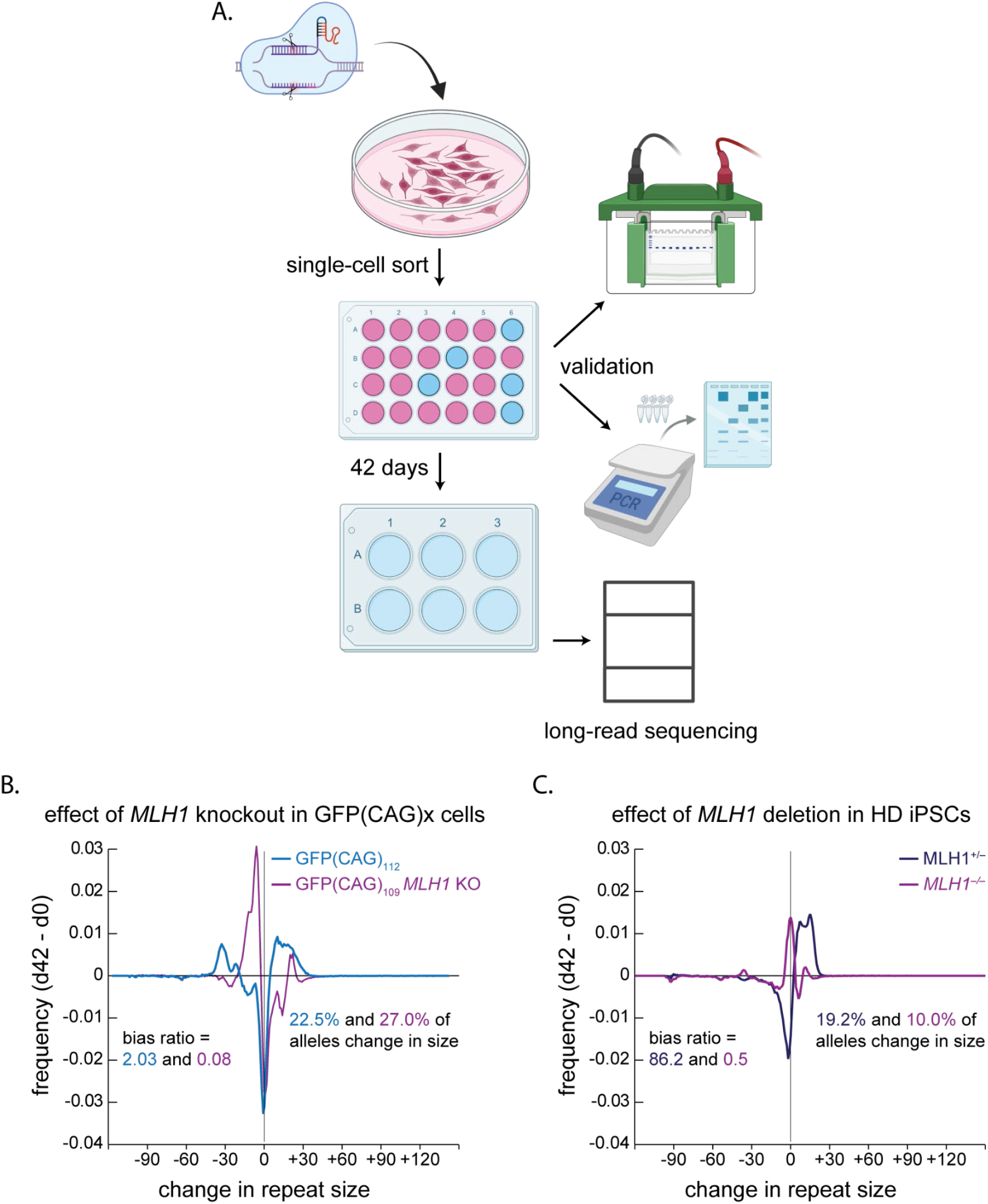
A quick protocol for the assessment of repeat instability phenotypes. A) Schematic of the work flow used for GFP(CAG)x cells. Cas9 RNPs are transfected into a cell population and the cells are sorted (or diluted) at a density of 1 cell per well. While they grow over the 42 day period, the cells must be split and the surplus cells can be used for western blotting and DNA analysis to confirm the knockout in each clone. At the end of the 42 days, the DNA from each clone is isolated and sequenced as before. B) Knocking out MLH1 on GFP(CAG)_109_ cells led to small contractions, but did not decrease the frequency of repeat expansions. C) Knocking out MLH1 in iPSCs led to a decrease in the frequency of instability, bias ratio, and the size of the contractions. Of note, the experimental flow is slightly different to accommodate iPSC-specific requirements (see methods).

Using this approach, we knocked out *MLH1* (Supplementary Figure 4), a treatment known to stabilize repeat tracts in mice and in cells (Iyer and Pluciennik 2021). Using SCIA, we found a significantly different delta-plot distribution compared to the parental line (P< 0.0001 - Figure 5b, Table 1). However, the instability frequency was 27.0%, which was similar to the wild-type parent (22.3%). The difference came from the marked bias towards contractions (bias ratio = 0.08). We conclude that this sped up protocol uncovers the expected effects of *MLH1* in repeat expansion and that SCIA coupled to delta-plot visualization led to a more accurate view of the effects engendered by the knockout.

### SCIA is applicable to HD iPSC-derived cells

Because HEK293-derived cells are not representative of any particular disease state or endogenous locus, we aimed to determine whether SCIA could work in HD-derived iPSCs. We therefore repeated SCIA using the same sgRNAs but this time nucleofected into iPSCs that contained a heterozygous expansion with 19 CAGs on one allele and 133 on the other. We adjusted the protocol to make single clones but to keep the cultures in larger plates so as to keep the cells pluripotent (See methods). Otherwise the same protocol was applied. We found similar results compared to the GFP(CAG)_116_ cells, in that the bias ratio was reduced, from 86.2 in the MLH1^+/–^ cells compared to 0.83 in the MLH1^−/−^ cells. However, in the HD iPSCs *MLH1* knockout also reduced the frequency of instability and the size of the changes (Table 2). We conclude that the SCIA protocol is compatible with iPSCs, making it applicable to multiple replicating cellular systems.

### Direct comparison between the instability index and SCIA

We next aimed to directly compare the ability of SCIA to find biologically relevant changes compared to the instability index. We therefore calculated instability indices from the same data obtained by the SCIA pipeline (Table 3). Where SCIA measured statistically significant changes in conditions when they were expected, the instability indices found significant changes in only one *MLH1* knockout clone in GFP(CAG)x-derived cells (P=0.05). These results suggest that instability indices can be less sensitive for some experimental designs in addition to providing fewer measures of instability.

**Table 3:** Instability indices for all lines used in this study.

| Cell type | Repeat size | Genotype | Treatment | Day | n= | Contraction index | Expansion index | Instability index‡ | P value** |
| --- | --- | --- | --- | --- | --- | --- | --- | --- | --- |
| GFP(CAG)x | 116 | WT | - | 0 | 1 | -7.69 | 1.89 | -5.81 | - |
|  |  | WT | Dox | 42 | 12 | -5.64 ± 4.0 | 4.61 ± 7.2 | -1.03 ± 9.1 | 0.10§ |
| GFP(CAG)x | 81 | WT | - | 0 | 1 | -4.56 | 0.91 | -3.65 | - |
|  |  | WT | Dox | 42 | 12 | -4.48 ± 4.2 | 1.65 ± 1.3 | -2.83 ± 5.3 | 0.86* |
| GFP(CAG)x | 110 | WT | - | 0 | 5 | -3.94 ± 0.48 | 3.71 ± 0.50 | -0.23 ± 0.97 | - |
|  |  | WT | - | 42 | 23 | -1.72 ± 2.0 | 6.81 ± 6.5 | 5.08 ± 7.7 | 0.10* |
| GFP(CAG)x | 118 | *FAN1* KO-1 | Dox | 0 | 1 | -3.86 | 2.30 | -1.56 | - |
|  |  | *FAN1* KO-1 | Dox | 42 | 11 | -5.26 ± 2.5 | 6.25 ± 3.5 | 0.99 ± 5.2 | 0.53* |
| GFP(CAG)x | 109 | *FAN1* KO-2 | Dox | 0 | 1 | -3.12 | 1.16 | -1.95 | - |
|  |  | *FAN1* KO-2 | Dox | 42 | 12 | -2.53 ± 0.72 | 5.31 ± 6.3 | 2.78 ± 6.4 | 0.35* |
| GFP(CAG)x | 108 | *PMS1* KO-1 | Dox | 0 | 1 | -2.35 | 0.71 | -1.64 | - |
|  |  | *PMS1* KO-1 | Dox | 42 | 7 | -3.26 ± 0.96 | 0.91 ± 0.22 | -2.35 ± 1.1 | 0.53* |
| GFP(CAG)x | 99 | *PMS1* KO-2 | Dox | 0 | 1 | -1.72 | 0.88 | -0.84 | - |
|  |  | *PMS1* KO-2 | Dox | 42 | 11 | -2.51 ± 0.56 | 0.75 ± 0.12 | -1.76 ± 0.68 | 0.64* |
|  | 109 | *MLH1* KO† | - | 0 | 1 | -3.35 | 4.26 | 0.90 | - |
|  |  | *MLH1* KO | - | 42 | 9 | -3.93 ± 3.8 | 3.55 ± 6.0 | -0.38 ± 8.7 | 0.05§ |
| HD iPSCs | 133 | *MLH1*<sup>+/+</sup> | - | 0 | 1 | -2.87 | 1.53 | -1.35 | - |
|  |  | *MLH1*<sup>+/-</sup> | - | 42 | 7 | -1.85 ± 0.74 | 3.65 ± 4.4 | 2.11 ± 5.1 | 0.094° |
|  |  | *MLH1*<sup>-/-</sup> | - | 42 | 14 | -2.21 ± 0.98 | 1.58 ± 2.0 | -0.646 ± 2.7 | 0.094° |

*WT* = parental wild-type. *dox* = doxycycline. \*: compared to *GFP(CAG)*<sub>116</sub> (+*dox*), §: compared to *GFP(CAG)*<sub>110</sub> (no *dox*), ‡: average ± standard deviation. All *P*-values are between expansion indices. \*\* FDR-corrected *P*-values between day 42 averages for instability indices only (see methods). †: This day 0 is a wild type population that was subsequently transfected with sgRNAs against *MLH1*. °: difference between HD iPSCs *MLH1*<sup>-/-</sup> and *MLH1*<sup>+/-</sup> expansion indices.

### Limitations of the SCIA quick protocol

We found, however, that we were limited by the efficiency of the Cas9 RNP. The Cas9 RNP against *MLH1* worked well as suggested by a PCR of the bulk population at sorting (Supplementary Figure 4 and Supplementary Figure 5) and we obtained 10 knockouts after screening 21 clones. However, our *FAN1* Cas9 RNPs were less effective, and we could not apply the optimised timeline since we obtained only 2 homozygous knockout clones after screening 53 clones. Thus, to fully take advantage of this protocol, highly effective Cas9 RNPs are essential.

## Discussion

We have provided a software to visualise the changes in repeat size distributions between two conditions without the need for complex coding on the part of the user. We showed that it can accommodate datasets from dividing and non-dividing cellular models as well as mouse tissues. One limitation is that it requires some treatment that engenders a change in repeat size (e.g., time) to be applicable. The analysis of the repeat size distribution of a single sample (e.g., one time point from one individual) would not benefit from the pipeline presented here. Comparing the instability index to SCIA analysis revealed that the instability index may be insensitive to changes perhaps because it does not take advantage of the wealth of data points that SCIA harnesses for statistical analysis. Importantly, the analysis pipeline presented here can differentiate between factors that have an effect on instability frequencies and those that have an effect on the size of the changes, which cannot be determined using the instability index.

The models of repeat instability always involve an initial trigger, which have been assumed to be caused by mismatch repair factors that can create a nick. This then creates the opportunity for error-prone repair that leads to a change in repeat size (Wheeler and Dion 2021). The prediction of this generalized model is that factors that increase the frequency of this initial trigger or the probability of error-prone repair should increase the instability frequency. By contrast, downstream processing factors that determine the size of the changes will see the frequency of processing unchanged but the size and/or bias ratio will be affected. We expect that some factors may work at multiple points in the pathway to repeat instability, affecting both frequency of instability and the size of the changes. This two step concept has often been overlooked when analysing repeat instability in cellular and animal models, but has been used in yeast [e.g., (Li et al. 2024). This is due in large part to the limitation of the assays used in mammals, which may only focus on changes of a certain size or direction [e.g., (Lin et al. 2006)] or be limited in resolving either very small (e.g., small-pool PCR) or very large changes (e.g., MiSeq sequencing). Thus, although the field has identified many genetic factors involved in repeat instability (Wheeler and Dion 2021), basic information about whether these factors increase the frequency of repeat size changes and/or affect the size of the changes are not well documented.

A significant insight gained here is that repeat tracts of 81 repeats have a frequency of repeat instability that is similar to those with 116 repeats, but the size of the changes is smaller when the repeat tract is lower. It is not yet clear whether this phenomenon extends to a wider range of repeat sizes or to other cell types. We also found that knocking out *MLH1* or *PMS1* had little effect on the frequency of repeat instability in GFP(CAG)_116_ cells, but the mutants had altered bias ratios and a reduced size of the changed alleles. One interpretation of this phenotype is that MLH1 and PMS1 affect the processing downstream of the initial trigger, at least in HEK293-derived cells. This is in contrast to the role of MLH1 in biochemical assays (Pluciennik et al. 2013; Senoussi et al. 2025). An alternative interpretation is that knocking out *MLH1* or *PMS1* in these cells lead to a shift in the balance between error-free and error-prone repair, perhaps via recruitment of FAN1 (Goold et al. 2021, 2019; Porro et al. 2021). Indeed, we found that *FAN1* affected the instability frequency while reducing the size of the expansions slightly. To complicate the matter, repeat instability is notoriously cell type specific (Dion 2014; Handsaker et al. 2025; Mätlik et al. 2024). Indeed, when we applied SCIA to knock out MLH1 in HD iPSCs, we found a decrease in the frequency of instability and in the size of the contractions. Moreover, the bias ratio skewed towards contractions. This is consistent with a cell-type specific role of repeat instability modifiers that will be worth investigating further.

We have developed a novel analysis pipeline that can be applied to non-dividing and dividing cells as well as mouse tissues. It is specifically designed to assess changes in repeat size over time or for comparing treatment. The experimental design and analysis pipeline presented here will help speed up the understanding of repeat instability mechanisms and the identification of drug targets that can ultimately provide much needed therapeutic avenues for expanded repeat disorders.

## Methods

### Repeat Detector

To determine repeat size, we used Repeat Detector (Taylor et al. 2022), which uses unaligned reads together with restrictive profiles with a repeat size range of [0–250]. For each analysis, the –with-revcomp option was enabled and data were output to a density plot (-o histogram option). The code for Repeat Detector can be found here: https://github.com/DionLab/RepeatDetector. Because of the presence of incomplete PCR fragments and short CAG stretches outside of the repeat tract, repeat sizes from 0 to 5 are removed in the downstream analyses.

### Delta plots, instability frequencies, bias ratios, and the average size of the changes

For the delta plots, the histograms of the number of CCSs per repeat size are normalized to the total number of CCS in each sample. Then we realigned the data to the mode at day 0 and plotted the change in repeat size. This is essential if the shape of the starting repeat size distribution is different, for example when comparing two different lines. We have done this in all cases, however, when the starting population is the same (e.g., when comparing the same line with and without doxycycline), this step can be skipped.

The area under the curve was calculated from the delta plots of the average of the samples at day 42 compared to their common distribution at day 0 (Figure 1ab). This is a measure of the instability frequency. All the frequencies above 0 on the delta plot were added together to make the instability frequency. The bias ratio was calculated as the ratio of the area under the curve for expansions divided by that of the contractions. The averages of size changes reported were calculated from the delta plots, using the average repeat sizes in the area under the curve for expansions and contractions separately.

### Analysis software

The GUI was developed using Python (version 3.13.5) together with wxPython library (version 4.2.3). Pandas (version 2.2.3) and NumPy (version 2.1.3) were used for data handling. Seaborn (version 0.13.2), with matplotlib (version 3.10.8) were used for visualization. RD is containerized within Docker (version 28.4.0) with .tar extension.

We tested the application on macOS (Darwin Kernel Version 24.6.0), and it is made to run on both Intel and latest Apple Silicon chip architectures. There is a singularity containerized object available to run on a HPC setting. Note that some of the scripts used here were coded with the help of AI.

The workflow of the software (version 2.0) is found in Supplementary Figure 1, and a working video example is also available (Supplementary Video 1). A detailed instruction is available on GitHub (https://github.com/DionLab/SCIA). The prerequisites are that the Docker desktop app is installed locally and running. The repeat-detector.tar file must be downloaded and present locally. The input for running RD and instability index is a FASTA file. FASTQ files were converted to FASTA using SeqKit. The user interface consists of two tabs. One tab is dedicated to Repeat Detector and instability indices. The user needs to specify which FASTA file is that of the control conditions. Both permissive and restrictive Repeat Detector profiles can be run, and a variety of output options are available to visualize repeat size distributions. The second tab uses the histogram outputs of Repeat Detector as input and generates the delta plots and runs the statistical analyses, including a KS test when two delta plots are compared. It returns a statistics summary containing the instability frequency, the average size of expansions and contractions, and the bias ratio.

### Publicly available and re-used long-read sequencing datasets

The HD iPSC-derived neurons have been generated before (Murillo et al. 2025) and they are available at https://www.ncbi.nlm.nih.gov/bioproject/PRJNA1077893. This dataset also contains the sequencing data from striatal samples of the HD mouse model R6/1 that were injected with AAVs expressing either the Cas9D10A alone or Cas9D10A together with a sgRNA against the repeat tract. We showed before that the presence of both AAVs led to contractions (Murillo et al. 2025).

### Cell lines and culture

HEK293-derived cells, GFP(CAG)x cells, where x is the number of repeats carried within the GFP-reporter inserted as a single copy on chromosome 12 (Ruiz Buendía et al. 2020; Santillan et al. 2014), and the knockout lines described here (Supplementary Table 2) were cultured as described (Larin et al. 2024; Cinesi et al. 2016; Mangin et al. 2024) in Dulbecco’s Modified Eagle’s Medium (DMEM) supplemented with 10% dialysed Foetal Bovine Serum (FBS), 1% Penicillin-Streptomycin, 15 μg ml^−1^ blasticidin and 150 μg ml^−1^ hygromycin. To initiate transcription through the GFP minigene, doxycycline was added to the assay media to a final concentration of 2 µg ml^-1^, except for the experiments using the *PMS1* knockout cells, which were done using 1 µg ml^-1^. The cultures were periodically tested for mycoplasma using the Eurofins service and found to be negative throughout this study.

### Induced pluripotent stem cells

HD iPSCs presented in Figure 1 were described before (HD iPSC Consortium 2012) and contain 19 CAG repeats on the non-expanded allele and 133 on the expanded allele. Cells were seeded in Corning Matrigel-coated plates and cultured under controlled conditions at 37°C, 5% CO_2_ in Gibco E8 Flex Medium, refreshed every 2 days. When passing, cells were washed with PBS, incubated with ReleSR (STEMCELL Technologies) for 5 minutes prior to re-plating on Matrigel-coated plates. For sample collection cells were detached from plates as described above, cell suspension was centrifuged at 300xg for 3 minutes and cell pellets stored at –80°C until DNA extraction.

### Single-clone isolation and instability assays

To obtain single cell clones, the starting culture was either serially diluted to obtain three 24-well plates to ensure a total of 8 to 12 colonies at the end of the experiment. Or, for some experiments, we used a BD FACSAria Fusion cell sorter to isolate single cells into 96-well plate formats. Both approaches have yielded similar results. An aliquot of the starting culture was collected to obtain day 0 DNA and/or proteins for comparison with later time points. The wells were monitored 4 to 7 days later and assessed for the presence of single colonies, with any wells containing >1 colonies being excluded. The media was replaced and cells were split as required. After 42 days, the cells were split into two pellets, one for DNA isolation using the Machery-Nagel NucleoSpin tissue-DNA extraction kit, one for protein extraction. For the experiments with *MLH1*, DNA and proteins were collected when the cells were split, such that the knockout and mutations could be confirmed as the clones grew.

For SCIA in iPSCs using *MLH1* sgRNAs, we used HD109 iPSCs, containing a stably integrated *NGN2* transgene at the CLYBL locus. This line was found to harbour 19 CAG repeats on the non-expanded allele and 133 repeats on the expanded one by SMRT sequencing. The cells were cultured on 6-well plates coated with Geltrex LDEV-Free Reduced Growth Factor Basement Membrane Matrix (diluted 1:100 in Knockout DMEM). Cultures were maintained at 37℃ in a humidified 5% CO2 incubator using Essential 8 (E8) Flex medium. Media was replenished every 2 days, with transgene selection maintained by supplementing media with 5μg/ml Blasticidin. Upon reaching ∼70% confluency, cells were washed with phosphate-buffered saline (PBS, without Ca^2+^ /Mg^2+^) and passaged twice weekly using ReLeSR enzyme-free selection reagent (STEMCELL technologies). To maintain cell viability and pluripotent colony architecture, cells were gently dissociated into small multicellular aggregates and reseeded at a split ratio between 1:8 to 1:14. Media was supplemented with 10μM ROCK inhibitor (Y-27632 dihydrochloride, Biotechne) for 24 hours immediately following thawing or single-cell plating to prevent apoptosis. Cellular morphology and confluency were regularly monitored via phase-contrast microscopy. The sgRNAs used were the same is in GFP(CAG)_116_ cells (Supplementary Table 2).

### CRISPR knockouts

For each target gene, two crRNAs (gRNAs) were designed to produce a 94 bp deletion (*FAN1 -* (McAllister et al. 2022)), a 68 bp deletion (*MLH1*), or a 97 bp deletion (*PMS1*) - See Supplementary Table 2). We used a tracrRNA labeled with ATTO-550 (IDT) for ease of sorting transfected cells. For each gene target, one million cells were nucleofected with Cas9 / crRNA / tracrRNA ribonucleoproteins (RNPs) using the 4D- Nucleofector and P3 Primary Cell 4D-Nucleofector X Kit and program CA137 (Lonza). After 24 hours, cells were sorted using a FACSAria Fusion to obtain the top 10% ATTO550 positive cells, which labels the tracrRNA, and were plated as single cells. The resulting clones were screened by PCR (see Supplementary Table 1 for primer sequences). We confirmed that the mutations were indeed present and determined the repeat size using Sanger sequencing or single -molecule real-time (SMRT) HIFI long-read sequencing (Supplementary Figures 2 (*FAN1* knockout), 3 (*PMS1* knockout), and 4 (*MLH1* knockout)).

To determine the editing efficiencies of the sgRNAs (Supplementary Figure 5), DNA was isolated from unsorted bulk populations, and the target site was PCR amplified. The band intensities for the edited and unedited bands were determined using ImageJ v1.54p. Band intensity was qualitatively proportional to the efficiency of sgRNA pairs, which was confirmed by counting the number of clones showing the predicted deletion in the sorted cells. Of note, the MLH1 sgRNAs were so efficient that we did not obtain wild type clones in this experiment (Supplementary Figure 6). However, MLH1 heterozygous knockout are expected to support normal levels of repeat expansion based on multiple cellular systems (Loupe et al. 2020; Pinto et al. 2013).

### Western blotting

For *FAN1, PMS1* and *MLH1* knockout lines, we also assessed protein expression by western blotting. Briefly, protein extraction was done using RIPA buffer containing EDTA-free Protease Inhibitor (Roche) as described (Murillo et al. 2025) with sonication (HeilsherUP50H, 1 second pulse, 60% amplitude). Proteins were quantified using the BCA Protein Assay kit (Pierce) as per manufacturers’ instructions. 30 µg of protein was loaded per lane of an 8-12% polyacrylamide gel and transferred onto a nitrocellulose membrane. The antibodies used and their concentrations are found in Supplementary Table 3. Fluorescence was visualised on a Licor Odyssey Clx scanner using Image Studio 6.1. All uncropped gels are found in Supplementary Figure 7.

### Long read targeted SMRT sequencing

Targeted sequencing of the repeat is carried out by PCR across the repeat locus and was done as described (Taylor et al. 2022; Larin et al. 2024). Briefly, we used the Mango Taq polymerase kit (Meridian Bioscience) with the primers listed in Supplementary Table 1. 50 µl PCR reactions were run using the following programme: 95°C for 5min followed by 32 cycles of 95°C for 30sec, 62°C for 30sec, and 72°C for 1min30 with a final extension of 72°C for 10min. PCRs were performed in triplicate and pooled, then purified with AMpure PB beads at a 0.6X volume ratio. 50 ng of purified PCR product per sample was prepared for sequencing using PacBio’s SMRTbell Express template preparation kit as per protocol number 101-791-700. Amplicons were barcoded using barcode adapter plate 3.0. Prepared amplicon libraries were quantified using Qubit HS dsDNA kit (ThermoFisher) and quality assessed using an Agilent fragment analyser HS NGS DNA kit. Between 10 and 12 pM of pooled libraries were loaded onto the SMRTcells and the sequencing was done using a Sequel IIe at Cardiff University School of Medicine. HIFI reads were used for the analysis as previously described (Taylor et al. 2022; Larin et al. 2024).

### Instability Indices

Instability indices were calculated as before (Murillo et al. 2025; Lee et al. 2010), using CCS frequency instead of the fluorescent signal used in the original study. So that background signal does not contribute to the instability index, the original study recommended setting a minimum threshold of signal intensity. Here we set this threshold at 5% of the frequency of the modal repeat size at day 0. However, because some sequencing datasets contain multiple peaks that can lead to incorrect results, we implemented a second threshold for the nearest repeat sizes on both sides of the mode that drops below 5% of the frequency of the modal repeat size. This avoids measuring a second allele that may be present, for example in patient samples. The instability index (II), contraction index (CI), and expansion index (EI) are calculated as:

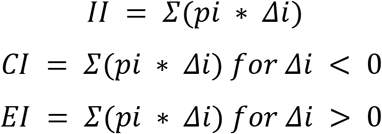

where pᵢ is the normalised peak height and Δᵢ is the index difference from the modal repeat size. When calculating the instability index for the HD iPSC samples, we only used repeat sizes above 30 CAGs to avoid interference by the wild type allele.

### Statistics

When comparing two delta plot distributions two-sample Kolmogorov–Smirnov (KS) tests were used. When comparing instability indices, we used two-tailed Mann-Whitney U tests between the indices obtained at day 42 for each clone. P-values were corrected for multiple comparisons using the Benjamini–Hochberg false discovery rate procedure, applied separately for each index. P≤0.05 was considered statistically significant throughout.

## Abbreviations

Dox: doxycycline
GUI: graphical user interface
KS test: Kolmogorov-Smirnov test
HD: Huntington’s disease
iPSC: induced pluripotent stem cells
SCIA: Single Clone-based Instability Assay (SCIA)
SMRT: PacBio Single Molecule Real Time (SMRT)

## Data Access

All the long-read sequencing dataset can be found on SRA (PRJNA1428164). The software and the source codes can be found at https://github.com/DionLab/SCIA, with the version controlled code available here: https://doi.org/10.5281/zenodo.18850398.

## Acknowledgements

We thank the members of the Dion lab for critical reading of the manuscript. Peter Holmans provided help with the statistics. We thank Nicholas Allen for the CRISPR knockout protocol and Thomas Massey for the *FAN1* and *MLH1* sgRNA sequences. The FAN1 antibody was a generous gift from CHDI. We would like to thank Joanne Morgan and the core team at the Centre for Neuropsychiatric Genetics and Genomics at Cardiff University for support with sequencing. Some of the figures were created using BioRender with (Licenses https://BioRender.com/oet71fi, https://BioRender.com/3hgcemc). This research was undertaken using the supercomputing facilities at Cardiff University operated by Advanced Research Computing at Cardiff (ARCCA) on behalf of the Cardiff Supercomputing Facility and the HPC Wales and Supercomputing Wales (SCW) projects. We acknowledge the support of the latter, which is part-funded by the European Regional Development Fund (ERDF) via the Welsh Government.

## Funding

This work was partly funded by Pfizer Inc, the UK Dementia Research Institute (UK DRI-3204) through UK DRI Ltd, principally funded by the UK Medical Research Council, and a grant from the Medical Research Council (MR/X02184X/1). The Dion lab is further supported by the Moondance Foundation Laboratory.

## Competing Interest Statement

V.D. declares that he has had a research contract with Pfizer Inc that made some of this work possible.

## Author contributions

CS: Conceptualization, Formal analysis, Investigation, Methodology, Visualization, Software, Writing – review & editing

RRPD: Formal analysis, Software, Writing – review & editing

ER: Conceptualization, Methodology, Investigation, Writing – review & editing ANA: Methodology, Investigation, Writing – review & editing

LH: Methodology, Writing – review & editing WE: Formal analysis, Writing – review & editing AM: Methodology, Writing – review & editing

VD: Conceptualization, Funding acquisition, Project administration, Supervision, Visualization, Writing – original draft, Writing – review & editing

## Declaration of generative AI and AI-assisted technologies in the manuscript preparation process

During the preparation of this work the authors used ChatGPT in order to assist with coding. After using this tool, the authors reviewed and edited the content as needed. They take full responsibility for the content of the published article.

## Supplementary Materials

**Supplementary Figure 1:**
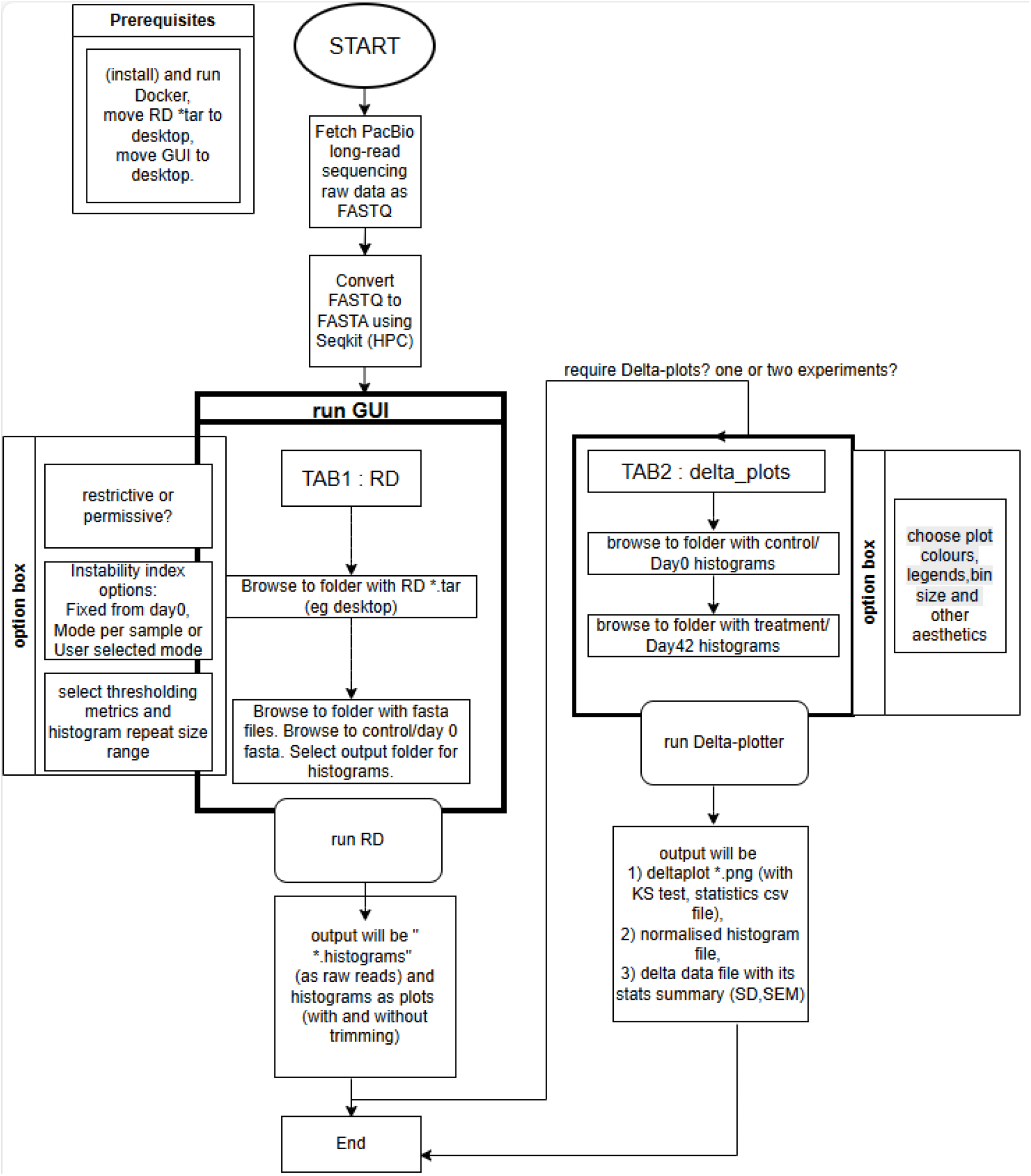
Flow chart of the software. RD = Repeat Detector. SD = standard deviation, SEM = standard error on the mean.

**Supplementary Figure 2:**
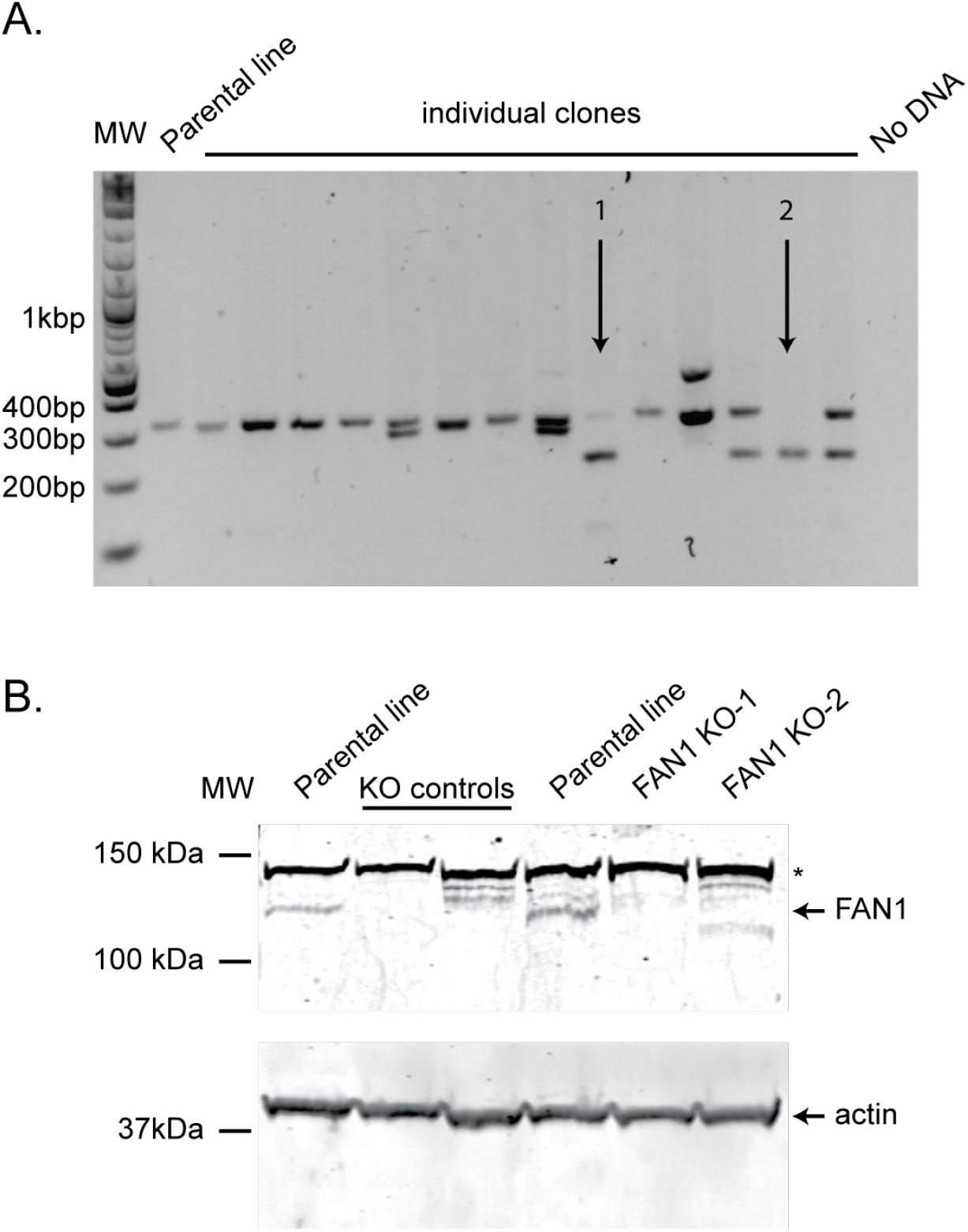
FAN1 knockout clone characterization in GFP(CAG)116. A) Individual clones after sorting were screened for the expected deletion of 94bp. Arrows show the clones that no longer had the wild type band. Note that clone 2 was a mixed clone and went through another round of single-cell sorting before being used here. The candidate clones were amplified, cloned and the mutations were sequenced using Sanger sequencing. B) Western blotting of the FAN1 knockout clones.

**Supplementary Figure 3:**
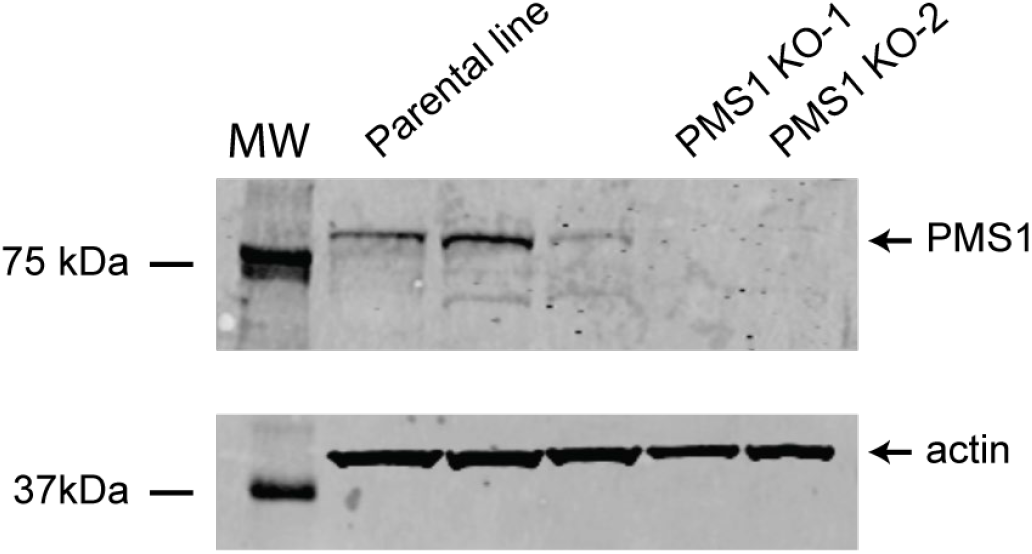
PMS1 knockout clone characterization in GFP(CAG)116. Western blotting of the PMS1 knockout clones.

**Supplementary Figure 4:**
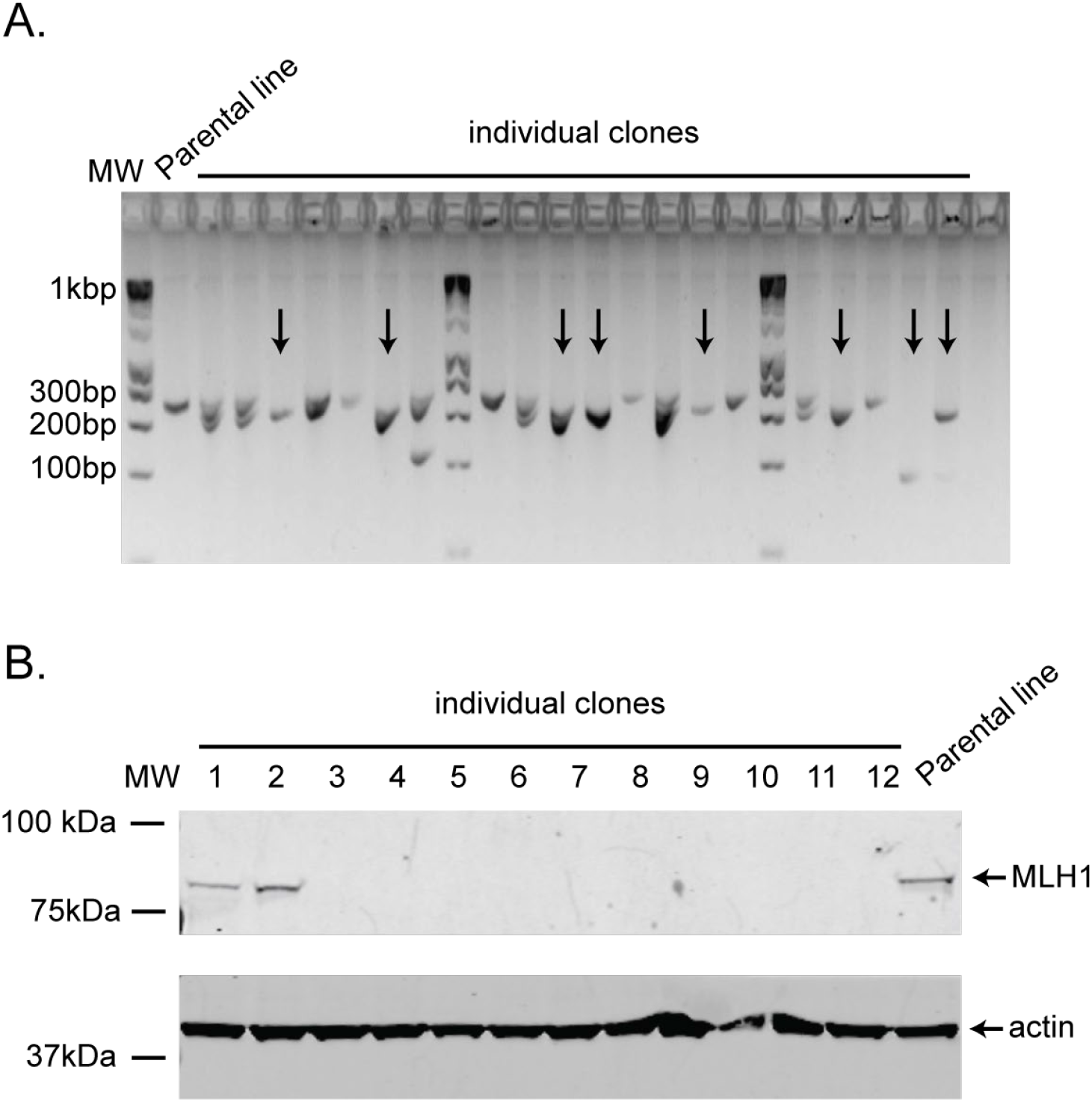
MLH1 knockout clone characterization in GFP(CAG)116. A) Individual clones after sorting were screened for the expected deletion of 68 bp. Arrows show the clones that no longer had the wild type band. The candidate clones were amplified, cloned and the mutations were sequenced using Sanger sequencing. B) Western blotting of the MLH1 knockout clones. Note that the first two clones were not deemed to be knocked out and were excluded from the MLH1 knockout datasets.

**Supplementary Figure 5:**
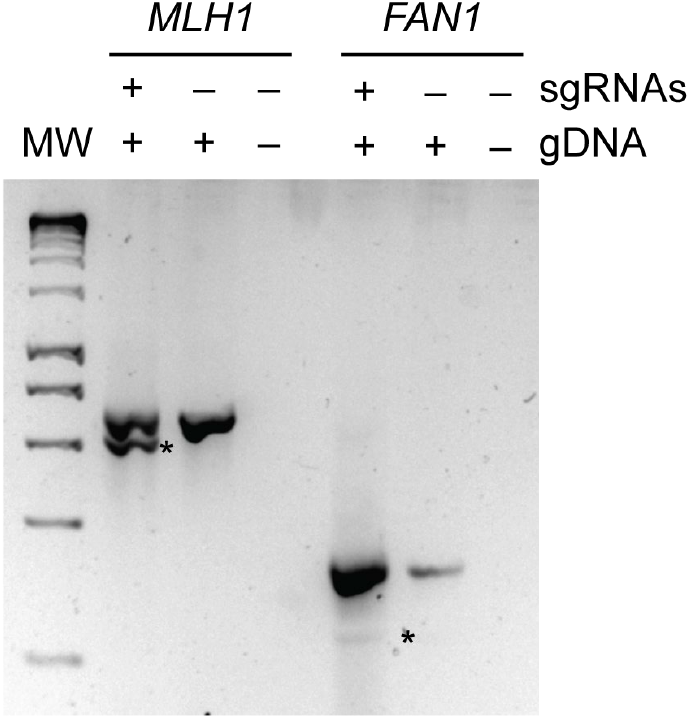
MLH1 and FAN1 sgRNA efficiencies in GFP(CAG)116. Gel showing the deletion expected by the use of the sgRNA pairs MLH1 and FAN1. The analyses were done on unsorted bulk populations. *: expected deletion product. gDNA: genomic DNA.

**Supplementary Figure 6:**
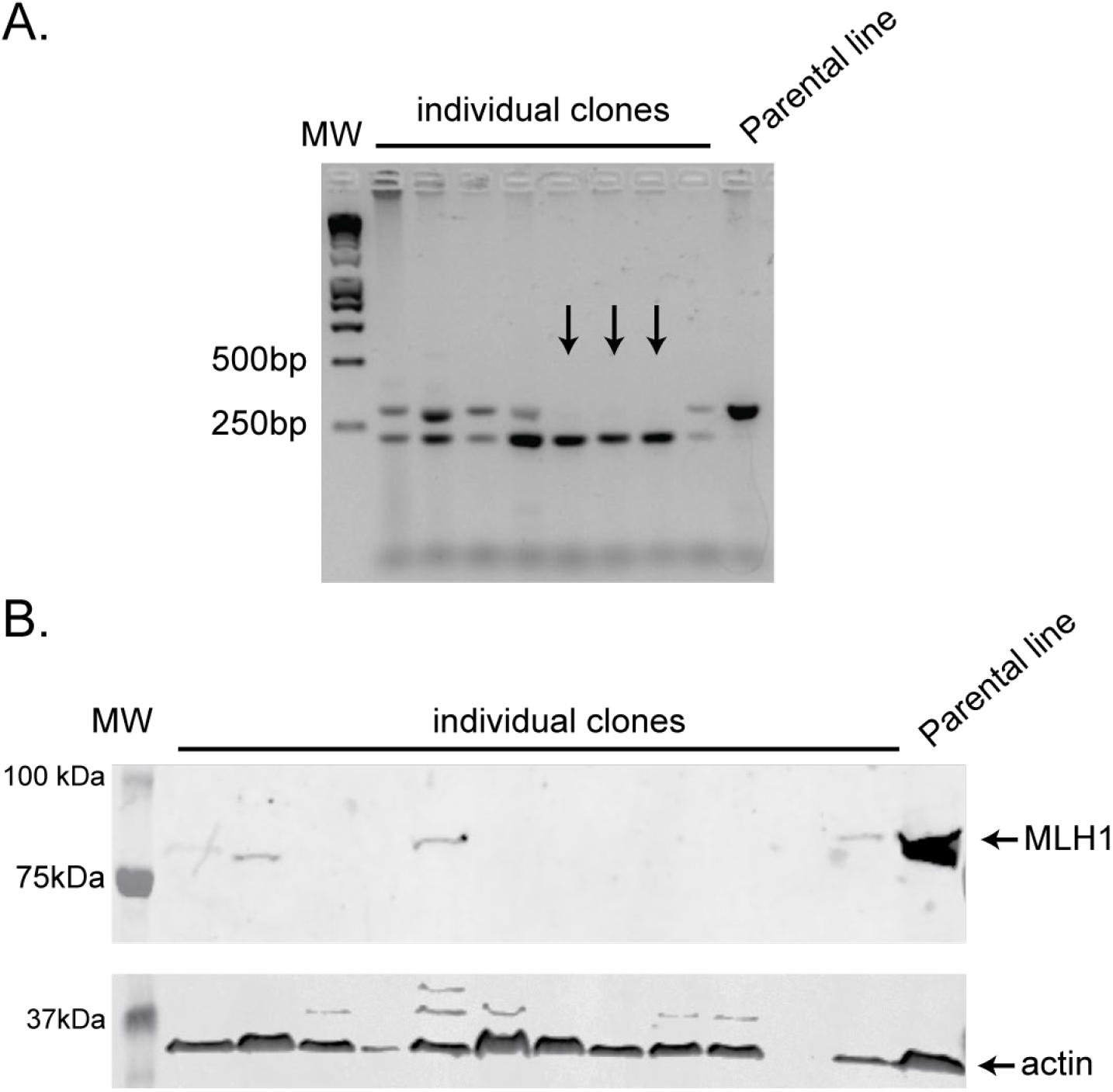
MLH1 knockout clones for SCIA in HD iPSCs. A) DNA gels showing a subset of the MLH1 genotyping in single HD iPSC clones after 42 days of culture. B) Western blot analysis example of some of the HD iPSC clones. Arrows indicate homozygous clones.

**Supplementary Figure 7:**
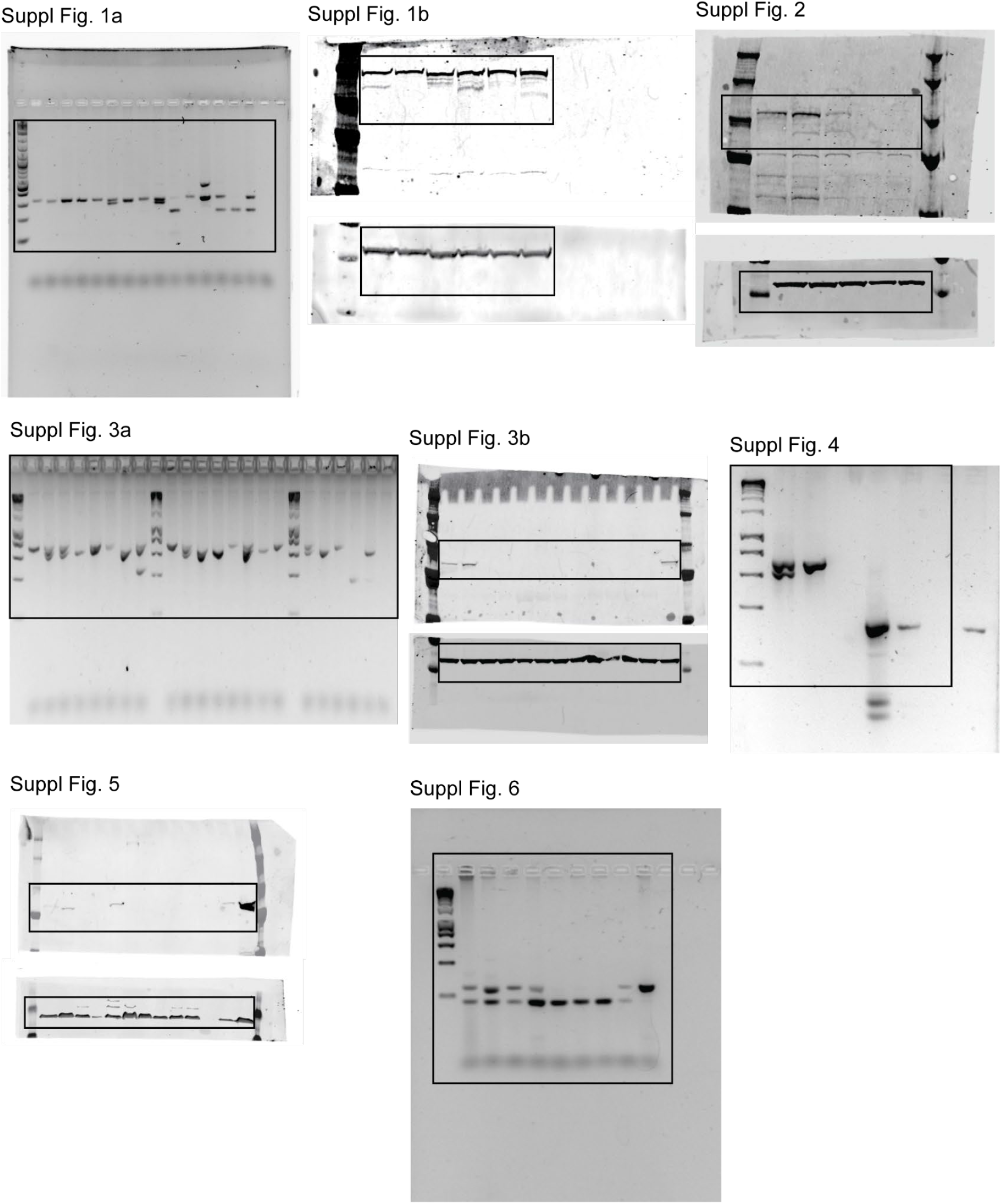
Original, uncropped gels and westerns. Black boxes show approximate cropping used in the figures.

**Supplementary Table 1:**
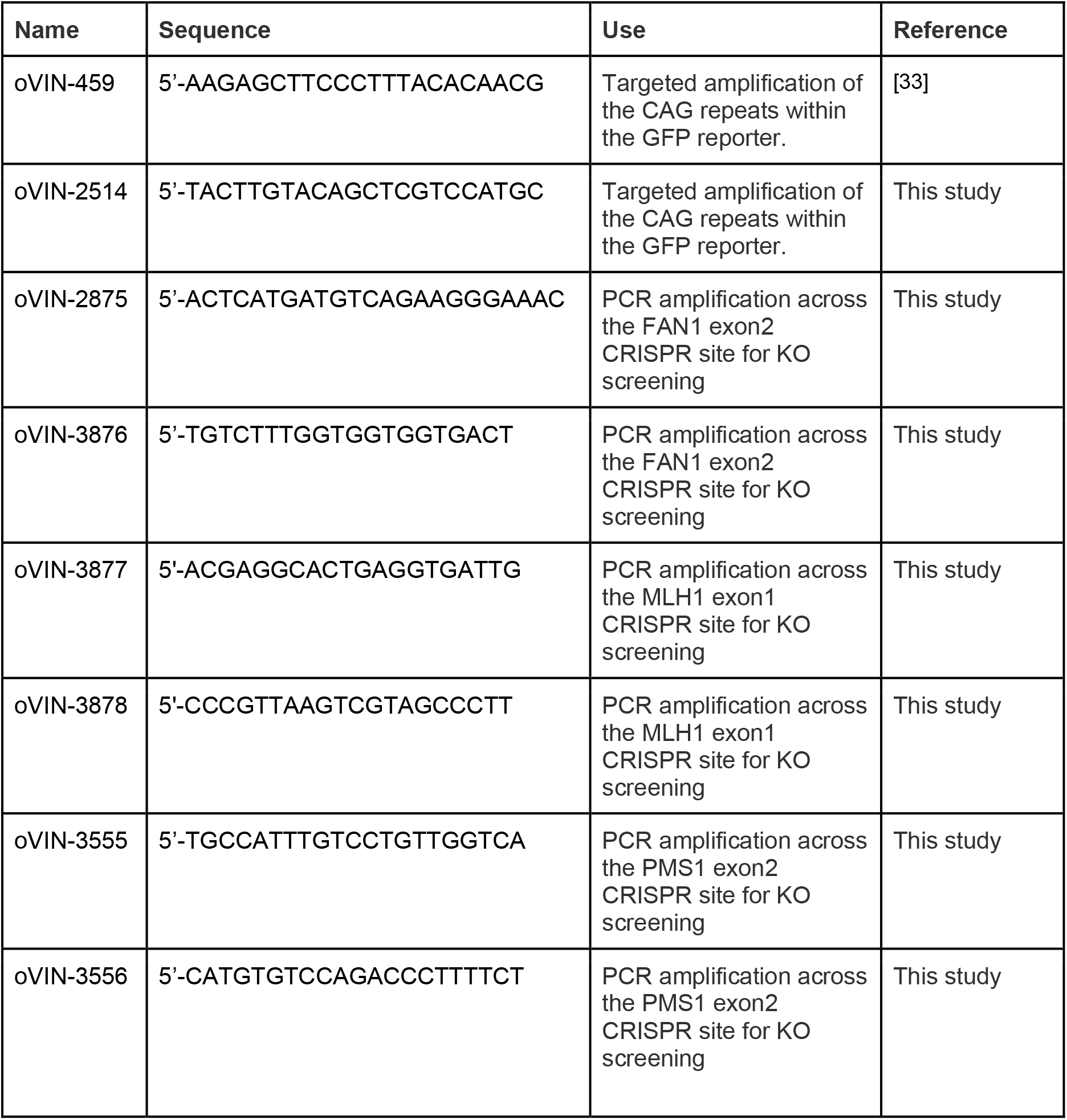
Sequence of primers used.

**Supplementary Table 2:**
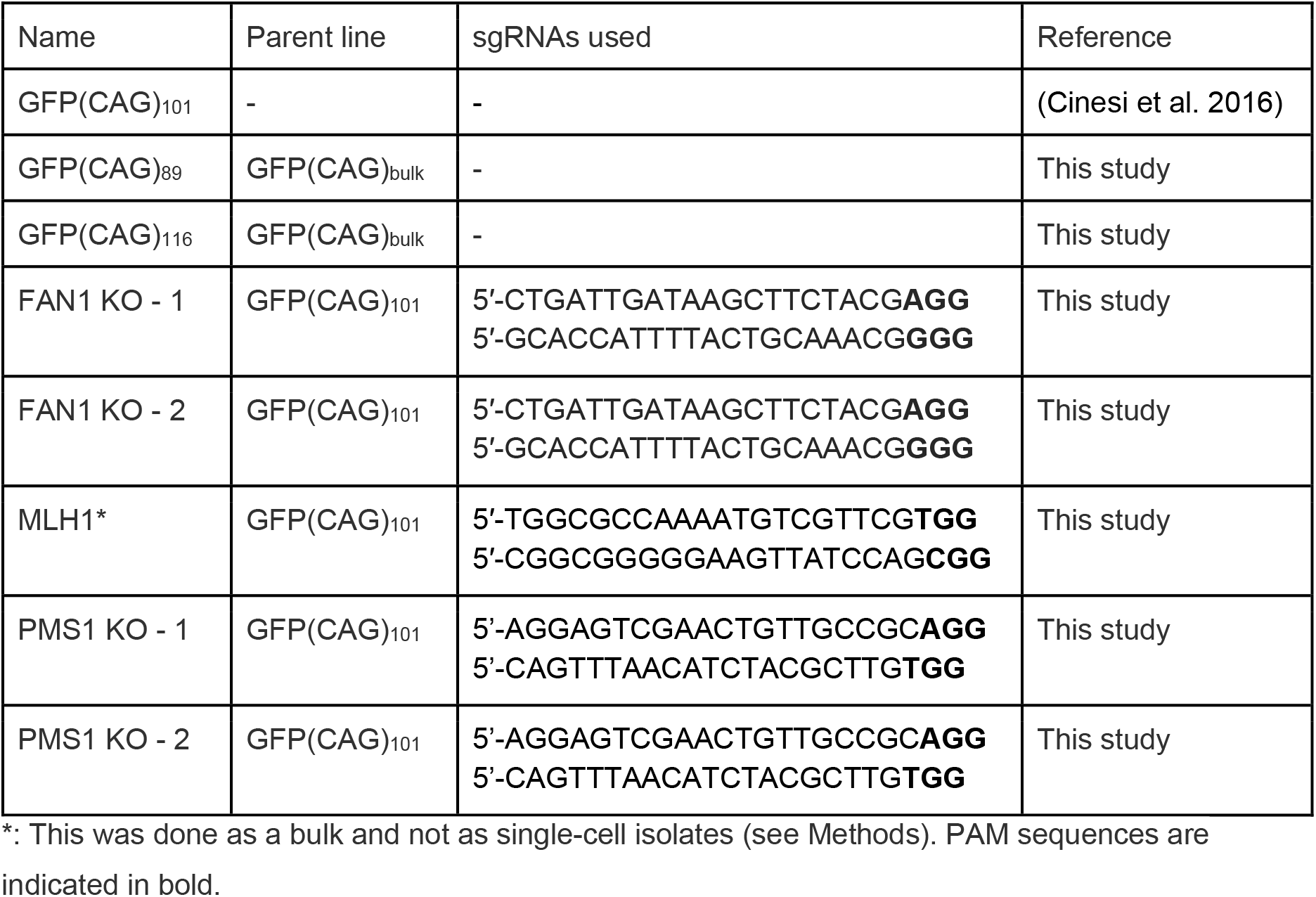
Cell lines used herein.

**Supplementary Table 3:** List of antibodies used.

| Antibody | Source | Species | Clonality | Concentration |
| --- | --- | --- | --- | --- |
| Anti-MLH1 | Abcam<br>#ab223844 | Rabbit IgG | Monoclonal | 1:1000 |
| Anti-FAN1 | CHDI | Sheep IgG | Polyclonal | 1:500 |
| Anti-β-Actin | Sigma #A5441 | Mouse IgG | Monoclonal | 1:15000 |
| Anti-PMS1 | Invitrogen PA5-35952 | Rabbit IgG | Polyclonal | 1:1000 |
| Donkey anti-Goat IgG (H+L) , Alexa Fluor™ 680 | Invitrogen A-21084 | Donkey IgG | Polyclonal | 1:10,000 |
| Goat anti-Mouse IgG (H+L) Alexa Fluor™ Plus 800 | Invitrogen A-21109 | Goat IgG | Polyclonal | 1:10,000 |
| Goat anti-Rabbit IgG (H+L) Alexa Fluor™ 680 | Invitrogen A32730 | Goat IgG | Polyclonal | 1:10,000 |

## Notes

https://doi.org/10.5281/zenodo.18850398

### Summary of Updates

We have added an important experiment where we show that the SCIA pipeline is applicable to a cellular Huntington's disease model, namely, induced-pluripotent stem cells.

## References

Bettencourt C, Hensman-Moss D, Flower M, Wiethoff S, Brice A, Goizet C, Stevanin G, Koutsis G, Karadima G, Panas M, et al. 2016. DNA repair pathways underlie a common genetic mechanism modulating onset in polyglutamine diseases. Annals of Neurology 79: 983–990.

Binda CS, Lelos MJ, Rosser AE, Massey TH. 2024. Chapter 13 - Using gene or cell therapies to treat Huntington’s disease. In Handbook of Clinical Neurology (eds. H.J. Federoff and R.A. Barker), Vol. 205 of Cell and Gene Therapies for Neurologic Diseases, pp. 193–215, Elsevier https://www.sciencedirect.com/science/article/pii/B9780323901208000149 (Accessed September 19, 2025).

Chen Z, Morris HR, Polke J, Wood NW, Gandhi S, Ryten M, Houlden H, Tucci A. 2024. Repeat expansion disorders. Practical Neurology. https://pn.bmj.com/content/early/2024/09/30/pn-2023-003938 (Accessed February 22, 2025).

Cinesi C, Aeschbach L, Yang B, Dion V. 2016. Contracting CAG/CTG repeats using the CRISPR-Cas9 nickase. Nat Commun 7: 13272.

Cleary JD, Pearson CE. 2005. Replication fork dynamics and dynamic mutations: the fork- shift model of repeat instability. Trends Genet 21: 272–280.

Deshmukh AL, Caron M-C, Mohiuddin M, Lanni S, Panigrahi GB, Khan M, Engchuan W, Shum N, Faruqui A, Wang P, et al. 2021a. FAN1 nuclease processes and pauses on disease-associated slipped-DNA repeats: Mechanism against repeat expansions. https://www.biorxiv.org/content/10.1101/2021.04.15.439995v1 (Accessed August 30, 2021).

Deshmukh AL, Porro A, Mohiuddin M, Lanni S, Panigrahi GB, Caron M-C, Masson J-Y, Sartori AA, Pearson CE. 2021b. FAN1, a DNA Repair Nuclease, as a Modifier of Repeat Expansion Disorders. J Huntingtons Dis 10: 95–122.

Dion V. 2014. Tissue specificity in DNA repair: lessons from trinucleotide repeat instability. Trends in Genetics 30: 220–229.

Donaldson J, Hensman Moss D, Ciosi M, Usdin K, Balmus G, Monckton DG, Tabrizi SJ. 2025. Huntington disease: somatic expansion, pathobiology and therapeutics. Nat Rev Neurol 1–17.

Ferguson R, Goold R, Coupland L, Flower M, Tabrizi SJ. 2024. Therapeutic validation of MMR-associated genetic modifiers in a human ex vivo model of Huntington disease. The American Journal of Human Genetics 111: 1165–1183.

Fernandez JP, Gärtner A, Perez-Perez J, Cumming SA, Feng J, Belois K van, He K, Yang D, Castells-Esteve C, Bergé-Gardeñes M, et al. 2026. Unscheduled DNA synthesis reveals a DNA repair hotspot and biomarker of somatic instability at the expanded-CAG repeat tract in the huntingtin gene. 2026.07.29.741355. https://www.biorxiv.org/content/10.64898/2026.07.29.741355v1 (Accessed August 19, 2026).

Genetic Modifiers of Huntington’s Disease (GeM-HD) Consortium. 2019. CAG Repeat Not Polyglutamine Length Determines Timing of Huntington’s Disease Onset. Cell 178: 887–900.e14.

Goold R, Flower M, Moss DH, Medway C, Wood-Kaczmar A, Andre R, Farshim P, Bates GP, Holmans P, Jones L, et al. 2019. FAN1 modifies Huntington’s disease progression by stabilizing the expanded HTT CAG repeat. Hum Mol Genet 28: 650–661.

Goold R, Hamilton J, Menneteau T, Flower M, Bunting EL, Aldous SG, Porro A, Vicente JR, Allen ND, Wilkinson H, et al. 2021. FAN1 controls mismatch repair complex assembly via MLH1 retention to stabilize CAG repeat expansion in Huntington’s disease. Cell Rep 36: 109649.

Handsaker RE, Kashin S, Reed NM, Tan S, Lee W-S, McDonald TM, Morris K, Kamitaki N, Mullally CD, Morakabati NR, et al. 2025. Long somatic DNA-repeat expansion drives neurodegeneration in Huntington’s disease. Cell 188: 623–639.e19.

Hayward B, Kumari D, Santra S, van Karnebeek CDM, van Kuilenburg ABP, Usdin K. 2024. All three MutL complexes are required for repeat expansion in a human stem cell model of CAG-repeat expansion mediated glutaminase deficiency. Sci Rep 14: 13772.

HD iPSC Consortium T. 2012. Induced Pluripotent Stem Cells from Patients with Huntington’s Disease Show CAG-Repeat-Expansion-Associated Phenotypes. Cell Stem Cell 11: 264–278.

Iyer RR, Pluciennik A. 2021. DNA Mismatch Repair and its Role in Huntington’s Disease. J Huntingtons Dis 10: 75–94.

Khristich AN, Mirkin SM. 2020. On the wrong DNA track: Molecular mechanisms of repeat-mediated genome instability. J Biol Chem 295: 4134–4170.

Kim JC, Harris ST, Dinter T, Shah KA, Mirkin SM. 2017. The role of break-induced replication in large-scale expansions of (CAG)n/(CTG)n repeats. Nat Struct Mol Biol 24: 55–60.

Larin M, Gidney F, Aeschbach L, Heraty L, Randall E, Heuchan AER, Buřičová M, Bérard M, Dion V. 2024. Cas9 Nickase-mediated contractions of CAG/CTG repeats are transcription-dependent and replication-independent. NAR Molecular Medicine ugae013.

Lea DE, Coulson CA. 1949. The distribution of the numbers of mutants in bacterial populations. Journ of Genetics 49: 264–285.

Lee J-M, Zhang J, Su AI, Walker JR, Wiltshire T, Kang K, Dragileva E, Gillis T, Lopez ET, Boily M-J, et al. 2010. A novel approach to investigate tissue-specific trinucleotide repeat instability. BMC Systems Biology 4: 29.

Li F, Phadte AS, Bhatia M, Barndt S, Monte Carlo Iii AR, Hou C-FD, Yang R, Strock S, Pluciennik A. 2025. Structural and molecular basis of PCNA-activated FAN1 nuclease function in DNA repair. Nat Commun 16: 4411.

Li L, Scott WS, Khristich AN, Armenia JF, Mirkin SM. 2024. Recurrent DNA nicks drive massive expansions of (GAA)n repeats. Proceedings of the National Academy of Sciences 121: e2413298121.

Lin Y, Dent SYR, Wilson JH, Wells RD, Napierala M. 2010. R loops stimulate genetic instability of CTG·CAG repeats. Proceedings of the National Academy of Sciences 107: 692–697.

Lin Y, Dion V, Wilson JH. 2006. Transcription promotes contraction of CAG repeat tracts in human cells. Nat Struct Mol Biol 13: 179–180.

Lin Y, Hubert L, Wilson JH. 2009. Transcription destabilizes triplet repeats. Mol Carcinog 48: 350–361.

López Castel A, Cleary JD, Pearson CE. 2010. Repeat instability as the basis for human diseases and as a potential target for therapy. Nat Rev Mol Cell Biol 11: 165–170.

Loupe JM, Pinto RM, Kim K-H, Gillis T, Mysore JS, Andrew MA, Kovalenko M, Murtha R, Seong I, Gusella JF, et al. 2020. Promotion of somatic CAG repeat expansion by Fan1 knock-out in Huntington’s disease knock-in mice is blocked by Mlh1 knock-out. Hum Mol Genet 29: 3044–3053.

Mangin A, Dion V, Menzies G. 2024. Developing small Cas9 hybrids using molecular modeling. Sci Rep 14: 17233.

Marzec P, Richer M, Lahue RS. 2025. Therapeutic targeting of mismatch repair proteins in triplet repeat expansion diseases. DNA Repair 147: 103817.

Massey T, McAllister B, Jones L. 2018. Methods for Assessing DNA Repair and Repeat Expansion in Huntington’s Disease. In Huntington’s Disease (eds. S.V. Precious, A.E. Rosser, and S.B. Dunnett), pp. 483–495, Springer, New York, NY 10.1007/978-1-4939-7825-0_22 (Accessed March 27, 2025).

Mathews EW, Coffey SR, Gärtner A, Belgrad J, Bragg RM, O’Reilly D, Cantle JP, McHugh C, Summers A, Fentz J, et al. 2025. Suppression of Huntington’s Disease Somatic Instability by Transcriptional Repression and Direct CAG Repeat Binding. Nat Commun 16: 10009.

Mätlik K, Baffuto M, Kus L, Deshmukh AL, Davis DA, Paul MR, Carroll TS, Caron M-C, Masson J-Y, Pearson CE, et al. 2024. Cell-type-specific CAG repeat expansions and toxicity of mutant Huntingtin in human striatum and cerebellum. Nat Genet 1–12.

McAllister B, Donaldson J, Binda CS, Powell S, Chughtai U, Edwards G, Stone J, Lobanov S, Elliston L, Schuhmacher L-N, et al. 2022. Exome sequencing of individuals with Huntington’s disease implicates FAN1 nuclease activity in slowing CAG expansion and disease onset. Nat Neurosci 25: 446–457.

McLean ZL, Gao D, Correia K, Roy JCL, Shibata S, Farnum IN, Valdepenas-Mellor Z, Kovalenko M, Rapuru M, Morini E, et al. 2024. Splice modulators target PMS1 to reduce somatic expansion of the Huntington’s disease-associated CAG repeat. Nat Commun 15: 3182.

Miller CJ, Kim G-Y, Zhao X, Usdin K. 2020. All three mammalian MutL complexes are required for repeat expansion in a mouse cell model of the Fragile X-related disorders. PLOS Genetics 16: e1008902.

Mirkin SM. 2007. Expandable DNA repeats and human disease. Nature 447: 932–940.

Mouro Pinto R, Murtha R, Azevedo A, Douglas C, Kovalenko M, Ulloa J, Crescenti S, Burch Z, Oliver E, Kesavan M, et al. 2025. In vivo CRISPR–Cas9 genome editing in mice identifies genetic modifiers of somatic CAG repeat instability in Huntington’s disease. Nat Genet 57: 314–322.

Murillo A, Alpaugh M, Durairaj RRP, Randall EL, Larin M, Heraty L, Aston AN, Taylor AS, Monteys AM, Stöberl N, et al. 2025. Cas9 Nickase-Mediated Contraction of CAG/CTG Repeats in Vivo is Accompanied by Improvements in Huntington’s Disease Pathology. 2024.02.19.580669. https://www.biorxiv.org/content/10.1101/2024.02.19.580669v2 (Accessed January 31, 2026).

Nakamori M, Pearson CE, Thornton CA. 2011. Bidirectional transcription stimulates expansion and contraction of expanded (CTG)•(CAG) repeats. Hum Mol Genet 20: 580– 588.

Phadte AS, Bhatia M, Ebert H, Abdullah H, Elrazaq EA, Komolov KE, Pluciennik A. 2023. FAN1 removes triplet repeat extrusions via a PCNA- and RFC-dependent mechanism. Proceedings of the National Academy of Sciences 120: e2302103120.

Pinto RM, Dragileva E, Kirby A, Lloret A, Lopez E, St Claire J, Panigrahi GB, Hou C, Holloway K, Gillis T, et al. 2013. Mismatch repair genes Mlh1 and Mlh3 modify CAG instability in Huntington’s disease mice: genome-wide and candidate approaches. PLoS Genet 9: e1003930.

Pluciennik A, Burdett V, Baitinger C, Iyer RR, Shi K, Modrich P. 2013. Extrahelical (CAG)/(CTG) triplet repeat elements support proliferating cell nuclear antigen loading and MutLα endonuclease activation. Proceedings of the National Academy of Sciences 110: 12277–12282.

Polleys EJ, House NCM, Freudenreich CH. 2017. Role of recombination and replication fork restart in repeat instability. DNA Repair (Amst) 56: 156–165.

Porro A, Mohiuddin M, Zurfluh C, Spegg V, Dai J, Iehl F, Ropars V, Collotta G, Fishwick KM, Mozaffari NL, et al. 2021. FAN1-MLH1 interaction affects repair of DNA interstrand cross- links and slipped-CAG/CTG repeats. Sci Adv 7: eabf7906.

Radchenko EA, McGinty RJ, Aksenova AY, Neil AJ, Mirkin SM. 2018. Quantitative Analysis of the Rates for Repeat-Mediated Genome Instability in a Yeast Experimental System. In Genome Instability: Methods and Protocols (eds. M. Muzi-Falconi and G.W. Brown), pp. 421–438, Springer, New York, NY 10.1007/978-1-4939-7306-4_29 (Accessed May 20, 2024).

Reddy K, Tam M, Bowater RP, Barber M, Tomlinson M, Nichol Edamura K, Wang Y-H, Pearson CE. 2011. Determinants of R-loop formation at convergent bidirectionally transcribed trinucleotide repeats. Nucleic Acids Res 39: 1749–1762.

Ruiz Buendía GA, Leleu M, Marzetta F, Vanzan L, Tan JY, Ythier V, Randall EL, Marques AC, Baubec T, Murr R, et al. 2020. Three-dimensional chromatin interactions remain stable upon CAG/CTG repeat expansion. Sci Adv 6: eaaz4012.

Santillan BA, Moye C, Mittelman D, Wilson JH. 2014. GFP-based fluorescence assay for CAG repeat instability in cultured human cells. PLoS One 9: e113952.

Senoussi I, Mengoli V, Cerana A, Rinaldi A, Marco A, Reginato G, Moro SG, Acharya A, Roy M, Jayachandran A, et al. 2025. Mechanism of trinucleotide repeat expansion by MutSβ- MutLγ and contraction by FAN1. Nat Commun 16: 9445.

Taylor AS, Barros D, Gobet N, Schuepbach T, McAllister B, Aeschbach L, Randall EL, Trofimenko E, Heuchan ER, Barszcz P, et al. 2022. Repeat Detector: versatile sizing of expanded tandem repeats and identification of interrupted alleles from targeted DNA sequencing. NAR Genomics and Bioinformatics 4: lqac089.

Veitch NJ, Ennis M, McAbney JP, Shelbourne PF, Monckton DG. 2007. Inherited CAG·CTG allele length is a major modifier of somatic mutation length variability in Huntington disease. DNA Repair 6: 789–796.

Wang N, Zhang S, Langfelder P, Ramanathan L, Gao F, Plascencia M, Vaca R, Gu X, Deng L, Dionisio LE, et al. 2025. Distinct mismatch-repair complex genes set neuronal CAG-repeat expansion rate to drive selective pathogenesis in HD mice. Cell 188: 1524–1544.e22.

Wheeler VC, Dion V. 2021. Modifiers of CAG/CTG Repeat Instability: Insights from Mammalian Models. J Huntingtons Dis 10: 123–148.

Yang B, Borgeaud AC, Buřičová M, Aeschbach L, Rodríguez-Lima O, Ruiz Buendía GA, Cinesi C, Taylor AS, Baubec T, Dion V. 2022. Expanded CAG/CTG repeats resist gene silencing mediated by targeted epigenome editing. Hum Mol Genet 31: 386–398.

Zhao X-N, Usdin K. 2018. FAN1 protects against repeat expansions in a Fragile X mouse model. DNA Repair (Amst) 69: 1–5.

